# Nonlinear readout of spatial cues underlies robustness of asymmetric cell division

**DOI:** 10.1101/2023.11.21.568006

**Authors:** Nelio T.L. Rodrigues, Tom Bland, KangBo Ng, Nisha Hirani, Nathan W. Goehring

## Abstract

A key challenge in the development of an organism is to maintain robust phenotypic outcomes in the face of perturbation. Yet, how such robust outcomes are encoded by developmental networks remains poorly explored. Here we use the *C. elegans* zygote as a model to understand sources of developmental robustness during PAR polarity-dependent asymmetric cell division. By quantitatively linking alterations in protein dosage to phenotype in individual embryos, we show that spatial information in the zygote is read out in a highly nonlinear fashion and, as a result, phenotypes are highly canalized against substantial variation in input signals. Specifically, our data point towards an intrinsic robustness of the conserved PAR polarity network that renders polarity axis specification resistant to variations in both the strength of upstream symmetry-breaking cues and PAR protein dosage. At the same time, we find that downstream pathways involved in cell size and fate asymmetry are similarly robust to dosage-dependent changes in the local concentrations of PAR proteins, implying non-trivial complexity in translating PAR signals into pathway outputs. We propose that “quantitative decoupling” of symmetry-breaking, polarity, and asymmetric division modules acts to suppress the accumulation of error as embryos move along this developmental trajectory, thereby ensuring that asymmetric division is robust to perturbation. Such modular organization of developmental networks is likely to be a general mechanism to achieve robust developmental outcomes.

## Introduction

Developmental systems possess a remarkable ability to give rise to stable phenotypes in the face of perturbations including variation in gene expression, noise, environmental conditions, physical insult or constraints, or even mutational load. This has led to the notion that systems may have evolved to minimize variance in outputs, i.e. phenotypic traits, in the face of perturbations or variation in input signals, rendering them robust (Gibson and Wagner, 2000; Siegal and Bergman, 2002).

Various mechanisms have been proposed to account for canalization of phenotypic outcomes (Félix and Barkoulas, 2015; Hallgrimsson et al., 2019; Kitano, 2004). These include dedicated ‘extrinsic’ mechanisms that compensate for perturbations, such as the buffering activity of heat shock proteins (Casanueva et al., 2012; Rutherford and Lindquist, 1998), microRNAs (Ebert and Sharp, 2012), or compensatory cell rearrangements (Jelier et al., 2016; Schnabel et al., 2006). There are also pathway-intrinsic mechanisms that emerge from features of developmental networks such as redundancy, molecular feedback, or nonlinear reaction pathways (Whitacre, 2012). For example, non-linearities in reaction pathways can yield threshold-like behaviors that effectively canalize variable input parameters into similar developmental trajectories, thereby allowing them to converge upon similar outcomes (Barkoulas et al., 2013; Green et al., 2017). Put another way, nonlinear responses allow downstream developmental modules to be ‘decoupled’ from variation or fluctuations in upstream modules (Kitano, 2004). In the case of mutational or allelic variation, such mechanisms can yield highly nonlinear genotype-phenotype maps.

Here we use the first cell division of the nematode *C. elegans* embryo as a model for exploring how the robustness of early embryonic processes is encoded by developmental networks. Due to its reproducibility of development and available tools for genetic, mechanical and cell biological perturbation, and quantitative analysis, *C. elegans* has emerged as a powerful model system for quantitative exploration of developmental pathways, including asymmetric cell division (Delattre and Goehring, 2021; Lang and Munro, 2017; Rose and Gonczy, 2014).

In *C. elegans* and related species, the first cell division is nearly always asymmetric in both size and fate, the latter manifest as cell cycle asynchrony between daughter cells (Delattre and Goehring, 2021). Decades of research have yielded insight into the molecular pathways that underlie asymmetry of *C. elegans* zygote division from symmetry-breaking to cell fate specification (Rose and Gonczy, 2014). The precise magnitude of these asymmetries can vary between species, but the size asymmetry and cell cycle asynchrony for each species are highly reproducible (Schulze and Schierenberg, 2011; Valfort et al., 2018). Several studies have looked at the effects of loss of size and timing asymmetries, demonstrating that embryos can tolerate substantial variation, at least in part through compensatory behaviors in later steps (Choi et al., 2020; Jankele et al., 2021). Yet, how precision of division asymmetry is achieved, why it is important, and how robust division asymmetry is to perturbation are largely unexplored.

Division asymmetry in the *C. elegans* zygote is under control of the conserved PAR polarity network (Kemphues et al., 1988)(Figure 1A). The PAR network consolidates spatial information provided by symmetry-breaking cues and transduces it to the various processes that ultimately orchestrate the zygote’s asymmetric division (Rose and Gonczy, 2014). These include both asymmetric spindle positioning and unequal segregation of cell fate determinants. The PAR proteins consist of two antagonistic groups of membrane-associated proteins that define opposing anterior and posterior membrane domains (Goehring, 2014; Lang and Munro, 2017). The anterior PAR proteins (aPARs), PAR-3, PAR-6, PKC-3 and CDC-42, are initially segregated away from the nascent posterior by actomyosin-driven cortical flows (Goehring et al., 2011; Munro et al., 2004) and restrict membrane association of the opposing posterior (pPAR) proteins PAR-1, PAR-2, LGL-1, and CHIN-1 through their phosphorylation by PKC-3 (Beatty et al., 2010; Folkmann and Seydoux, 2019; Hao et al., 2006; Hoege et al., 2010). Conversely, pPAR proteins exclude aPARs through the phosphorylation of PAR-3 by PAR-1 (Benton and St Johnston, 2003; Motegi et al., 2011) and inhibition of active CDC-42 by the CDC-42 GAP CHIN-1 (Beatty et al., 2013; Kumfer et al., 2010; Sailer et al., 2015). At the same time, a second semi-redundant flow-independent pathway involving RING-dependent dimerization of PAR-2 and its local protection from PKC-3 activity by microtubules facilitates posterior pPAR membrane association (Bland et al., 2023; Motegi et al., 2011; Zonies et al., 2010). Once polarity is established, asymmetric PAR activity guides asymmetric positioning of the spindle and partitioning of cytoplasmic fate determinants (Rose and Gonczy, 2014).

**Figure 1.**
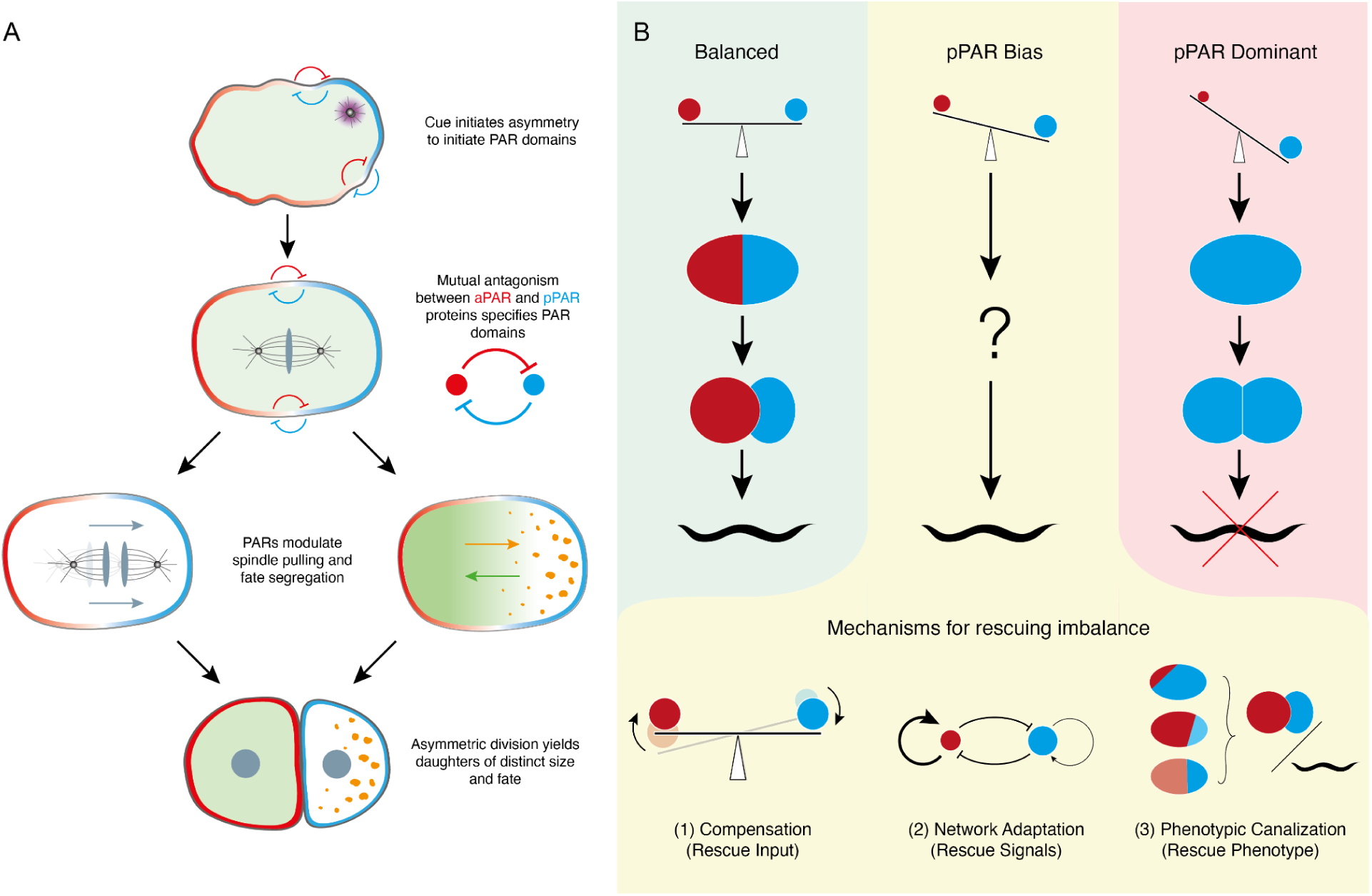
Potential mechanisms for dosage robustness in asymmetric division. **(A)** Schematic of the asymmetric division pathway in *C. elegans* zygotes. A local cue induces asymmetry of PAR proteins which is then reinforced by mutual antagonism between anterior and posterior PAR proteins (aPAR, pPAR) to generate stable domains. PAR proteins then spatially regulate downstream processes to drive division asymmetry. Due to this mutually antagonistic relationship, aPAR and pPAR protein levels/activities must be reasonably balanced to achieve proper polarity. **(B)** In wild-type embryos, relative levels of anterior and posterior PAR proteins are balanced to yield near equally sized domains and divisions are asymmetric (green, left). Strong depletion or homozygous mutation of a given PAR protein disrupts this balance, leading to dominance of one set of PAR proteins and failure in zygotic polarization, leading to symmetric division and loss of viability (red, right). While we know that embryos are generally robust to heterozygosity in *par* genes (yellow, middle), how polarity pathways compensate for modest imbalance in PAR dosage/activity - if they do at all - is unclear. We consider several potential models: (1) compensation - PAR protein levels actively adapt to restore balance; (2) network adaptation - features of the network compensate for imbalance to maintain stable polarity signals, such as changes in feedback strength or pattern; (3) signal canalization - downstream pathways are quantitatively robust to variability in polarity signals.

Polarization and asymmetric division of *C. elegans* zygotes is often described as robust (Lang and Munro, 2017; Motegi and Seydoux, 2013). Both processes are resilient to a number of genetic and environmental perturbations, including both temperature variation and physical deformation (Begasse et al., 2015; Klinkert et al., 2019; Labbé et al., 2006; Neves et al., 2015; Schenk et al., 2010). Polarization of the zygote still occurs in embryos with substantial defects in symmetry-breaking cues (Delattre and Goehring, 2021; Gross et al., 2019; Motegi et al., 2011; Zonies et al., 2010) and mispositioned or wrongly sized PAR domains can be corrected (Mittasch et al., 2018; Schenk et al., 2010). Finally, we know that *C. elegans* development is generally insensitive to heterozygous mutations in the majority of genes essential for early embryogenesis (Hodgkin, 2005), including all *par* genes (Kemphues et al., 1988; Watts et al., 1996). Despite the apparent robustness of polarity, there have been few systematic attempts to quantitatively link perturbations to division outcomes in this system (see examples in (Jankele et al., 2021; Pecreaux et al., 2006)).

PAR polarity is generally thought to require balance between the activities of aPARs and pPARs acting via the polarity kinases PKC-3 and PAR-1 (Goehring et al., 2011; Labbé et al., 2006; Lim et al., 2021; Watts et al., 1996), which in turn provide the key signals to spatially regulate downstream pathways (Galli et al., 2011; Griffin et al., 2011; Tenlen et al., 2008). Thus, models generally predict that polarization and asymmetric division of the zygote would be sensitive to PAR dosage changes. On its face, this prediction would seem to contradict the lack of obvious phenotypes in embryos heterozygous for *par* mutations, thus raising the fundamental questions of how sensitive this system really is to changes in PAR protein concentrations, and if it is indeed robust to dosage changes, what design features underlie this robustness.

To address these questions, we set out to explicitly measure the sensitivity of embryos to perturbations in PAR protein dosage. By combining established methods for manipulation of protein dosage in *C. elegans* (Oegema, 2006) with a recently developed image quantitation-based workflow (Rodrigues et al., 2022), we were able to directly relate dosage to phenotype in individual embryos. Our data support a model in which pathway responses are highly canalized against variation in spatial signals at multiple levels, leading to quantitative decoupling between symmetry-breaking, polarity, spindle positioning and fate segregation modules. This decoupling effectively prevents variability in a given module from propagating along this developmental trajectory and ensures highly reproducible outcomes in both size and fate asymmetry during asymmetric division.

## RESULTS

We reasoned that robustness of developmental processes to changes in gene or protein dosage in a given molecular network could arise from: (1) *Dosage compensation*, in which animals harboring a loss of function allele in a given locus could simply upregulate expression of the remaining functional allele, ensuring normal concentrations. Alternatively, there could be compensatory changes to levels of other molecules within the network to restore function to normal. (2) *Network adaption*, by which feedback pathways would allow the relevant network to adapt to dosage changes and thus function similarly to the wild type condition with respect to input and output signals. (3) *Phenotypic canalization* in which the network behavior itself is dosage sensitive, but downstream pathways are insensitive to variation in the network’s output signals (Figure 1B).

### Compensatory dosage regulation cannot explain robustness to heterozygosity in *par* **genes.**

As a first step towards distinguishing between these potential models, we asked whether embryos exhibited dosage compensation. Chromosome-wide dosage compensation is well known in the context of sex chromosomes: gene expression is systematically up- or down-regulated to account for differences in sex chromosome number in males and females (Jordan et al., 2019). However, dosage compensation of individual autosomal genes is less well understood. Systematic transcriptional analysis suggests the degree of compensation can vary substantially at the level of individual genes, though the vast majority show no or only partial compensation (Malone et al., 2012; Ragipani et al., 2022).

We initially looked for evidence of dosage compensation in animals heterozygous for mutant alleles of two polarity genes *par-6* and *par-2*, as representatives of aPAR and pPAR proteins, respectively. Due to maternal provision to oocytes, the mRNA and protein composition is primarily determined by the mother’s genotype. Thus, for simplicity, hereafter we refer to embryos by the genotype of the mother, i.e. heterozygous embryos = embryos from heterozygous mothers. To test whether compensation occurs, we applied spectral autofluorescence correction using SAIBR (Rodrigues et al., 2022) to accurately quantify and compare GFP levels in embryos of three genotypes: (1) homozygous for endogenously *gfp*-tagged alleles (*gfp/gfp*) in which all protein is GFP-tagged; (2) heterozygous embryos carrying a single tagged allele together with an untagged wild-type allele (*gfp/+*), in which we expect GFP-labeled protein to constitute roughly half of total protein; and (3) heterozygous embryos carrying a single tagged allele over either a null allele or an allele that can be selectively depleted by RNAi (*gfp/-*). For perfect dosage compensation, we would expect levels of GFP in *gfp/gfp* and *gfp/-* to be similar (Figure 2A). However, we find that embryos from *gfp/-* worms expressed levels of GFP that were only modestly increased relative to *gfp/+*, and well below those of *gfp/gfp* embryos, suggesting only partial compensatory upregulation (Figure 2B, 2C, 2D, Supplemental Figure S1). Similar results were obtained for other *par* genes examined, including *par-1*, *par-3*, and *pkc-3*, which showed modest to no compensation in heterozygotes (Figure 2D, Supplemental Figure S1).

**Figure 2.**
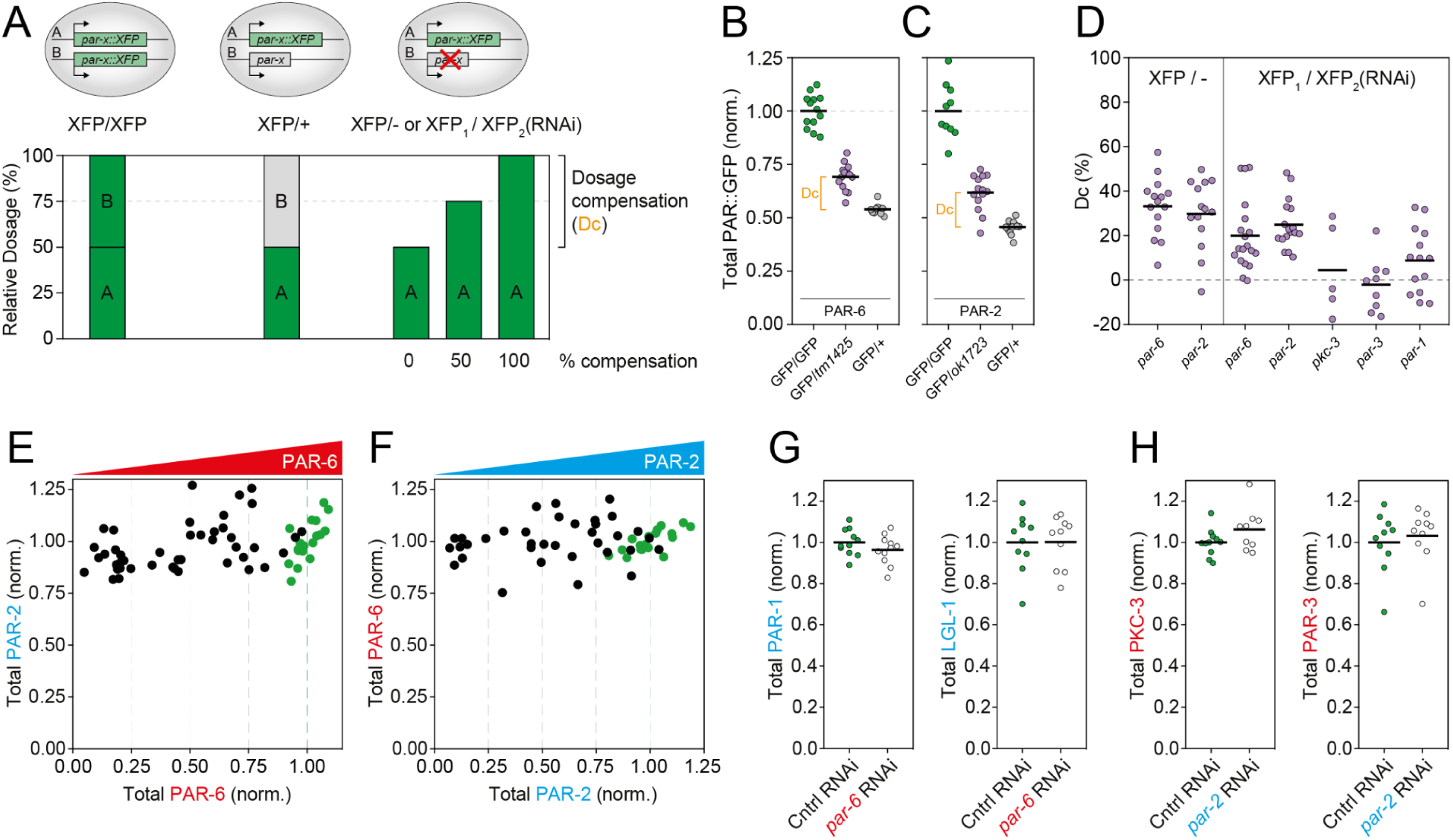
Minimal compensatory regulation in response to *par* gene / protein dosage changes. **(A)** Schematic for dosage compensation assay. Levels of XFP (GFP or mNG) were measured for embryos of three genotypes: homozygous, carrying two copies of an *XFP::par* allele (*xfp/xfp*); heterozygous, carrying one copy of *XFP::par* allele and one untagged allele (*xfp/+*), which is expected to express XFP at ∼50% levels of homozygotes; and heterozygous, carrying one copy of the *XFP::par* allele and either a mutant or RNAi-silenced allele (*xfp/-* or *xfp/RNAi*). Dosage compensation is quantified as the degree of excess XFP signal in *xfp/-* or *xfp/RNAi* embryos, expressed as the fraction of the difference in XFP signal between *xfp/xfp* and *xfp/+* animals. **(B-C)** Normalized GFP concentrations of PAR-6::GFP (B) or GFP::PAR-2 (C) as measured in embryos with the indicated genotypes: homozygous (*gfp/gfp*), heterozygous mutant (*gfp/-*) and heterozygous untagged (*gfp/+*) genotypes. **(D)** Modest or no dosage compensation exhibited for various *par::XFP* gene fusions when expressed in a heterozygous condition together with either a mutant (XFP/-) or an RNAi-silenced allele (XFP_1_/XFP_2_(RNAi)). Additional details for allele-specific RNAi in Supplemental Figure S1. **(E)** Total PAR-2 concentration is constant as a function of PAR-6 dosage. Embryos expressing both mCh::PAR-2 and PAR-6::mNG (expressed from the endogenous loci) were subjected to progressive depletion of PAR-6 by RNAi and total concentrations of mNG and mCh measured. Green datapoints are embryos treated with control RNAi (ie, showing wild-type protein levels). **(F)** Total PAR-6 concentration is constant as a function of PAR-2 dosage. Fluorescence tags as in (A), but embryos were subjected to progressive depletion of PAR-2 by RNAi. Note similar results were obtained using reversed tags (GFP::PAR-2 and PAR-6::mCh; data not shown). **(G)** PAR-1 and LGL-1 concentrations are unchanged in *par-6(RNAi)*. **(H)** PKC-3 and PAR-3 concentrations are unchanged in *par-2(RNAi)*. In B-D, G, H, individual embryo values shown with mean indicated.

Because of the requirement for balanced activity of aPAR and pPAR proteins (Goehring, 2014; Lang and Munro, 2017), we also asked whether down-regulation of other components in the PAR network could help explain the robustness of embryos to dosage changes in individual PAR proteins. In other words, would depletion of a given PAR protein lead to reduction in the concentration of opposing PAR proteins? We found that dosage of PAR-2 remained constant across the full range of PAR-6 depletion conditions and that PAR-6 levels were similarly constant across the full range of PAR-2 depletion (Figure 2E, 2F). Consistent with these results, we found that the other posterior PAR proteins PAR-1 and LGL-1 were unchanged in PAR-6-depleted animals (Figure 2G), while the levels of anterior PAR proteins PKC-3 and PAR-3 were unchanged in PAR-2-depleted animals (Figure 2H). Thus, there do not appear to be coordinated alterations in protein amounts to compensate for changes in the dosage of a given PAR protein.

We conclude that *C. elegans* embryos do not exert homeostatic regulation of PAR concentrations in response to dosage changes. It is possible that modest up-regulation of protein amounts for some *par* genes (*par-1*, *par-2*, *par-6*) could partially contribute to stable phenotypes in heterozygotes. However, the limited degree of upregulation means that heterozygotes are viable despite harboring 30-50% less PAR protein than wild type, raising the question of how dosage variation impacts polarity and asymmetric division.

### Asymmetric division is robust to changes in PAR dosage

We next turned to determining how alterations in PAR dosage impacted phenotypes associated with asymmetric division. We focussed on two key outputs: daughter size asymmetry, which is controlled by asymmetric spindle positioning, and cell cycle asynchrony of the AB and P1 daughters, which is manifest as a roughly two minute cell cycle delay in division timing and is a commonly used proxy for the asymmetric partitioning of cytoplasmic fate determinants (Schematic in Figure 3A, 3B)(Deppe et al., 1978; Kemphues et al., 1988; Rivers et al., 2008; Sulston et al., 1983).

**Figure 3.**
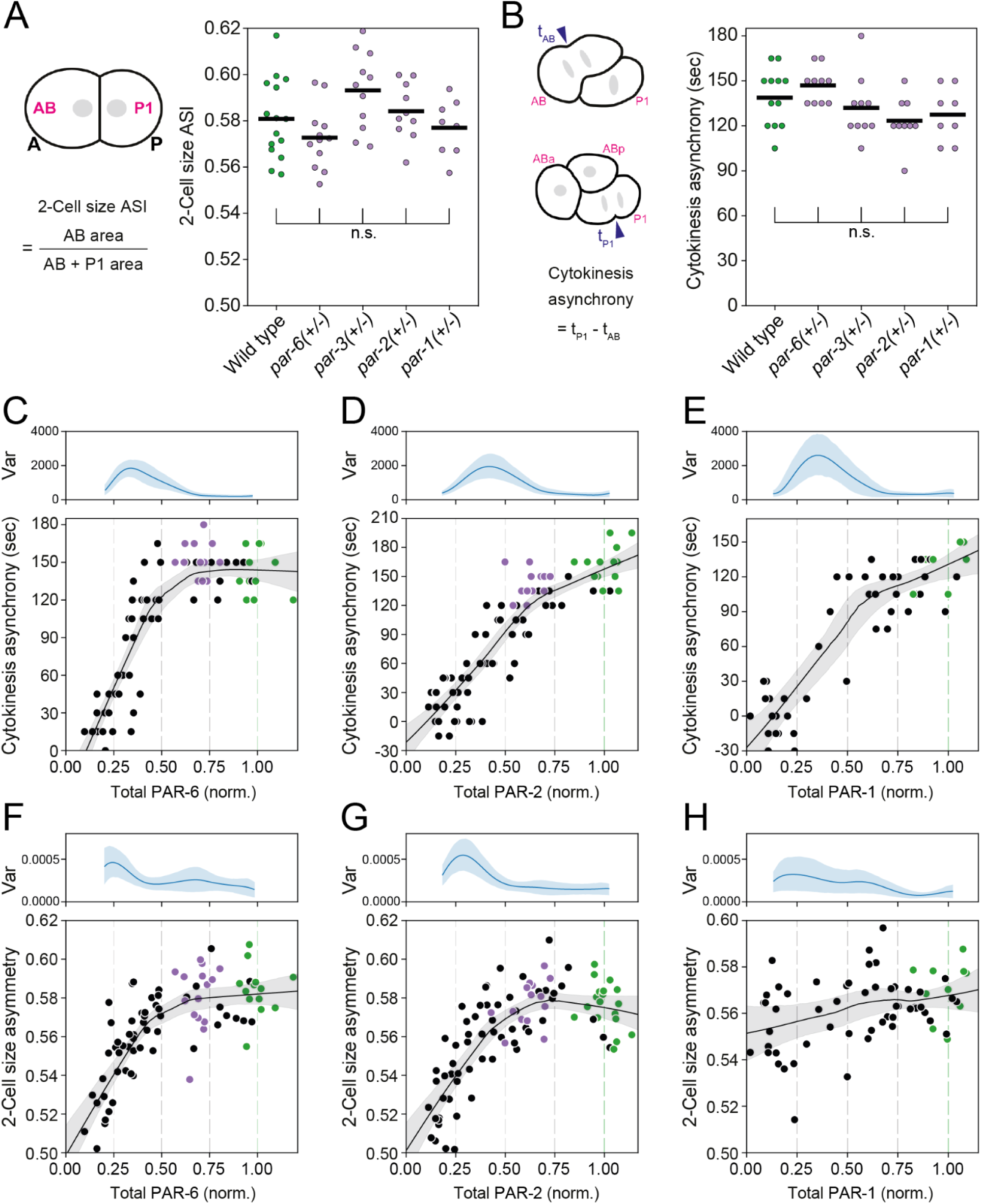
Division asymmetry is robust to changes in PAR dosage. **(A-B)** Size asymmetry (A) and asynchrony in cleavage furrow initiation (B) for AB and P1 blastomeres of the 2-cell embryo in embryos heterozygous for mutations in the *par* genes indicated. Genotypes: *+/+* (wild-type), *par-6(tm1425/+)*, *par-3(tm2716/+)*, *par-2(ok1723/+)*, and *par-1(tm2524/+)*. Heterozygous samples were compared to wild-type using an unpaired t test, where n.s. indicates no statistical difference. **(C-H)** AB vs P1 asynchrony (C-E) and size asymmetry (F-H) as a function of total dosage of PAR-6 (C,F), PAR-2 (D,G), and PAR-1 (E,H). Data for individual embryos subject to RNAi shown (black), compared with embryos from wild-type control (green) and in the case of PAR-2 and PAR-6, heterozygous animals (*gfp/-*, purple). Lines indicate Lowess smoothing fit with 95% confidence interval determined by bootstrapping to help visualize trends. Phenotypic variance (Var, see Methods) as a function of dosage is indicated above each panel.

To broadly determine how asymmetry depends on PAR protein dosage, we scored relative size asymmetry and division time asynchrony in embryos heterozygous for *par-1, par-2*, *par-3*, and *par-6* (Figure 3A, 3B). In all heterozygotes, we observed only minor statistically non-significant changes in size asymmetry and division asynchrony that were within the standard deviation observed in wild-type embryos. Thus heterozygotes are capable of robustly achieving both size and fate asymmetry despite changes in PAR dosage.

We then used progressive depletion of PAR proteins by RNAi to quantify the relationship between protein dosage and division asymmetry. Plots of dosage vs timing asynchrony for embryos depleted for PAR-1, PAR-2 or PAR-6 were generally nonlinear, with inflection points located at or near the point of 50% depletion (Figure 3C-E). Similarly, for all three proteins analyzed, (PAR-1, PAR-2, and PAR-6) size asymmetry remained within the wild-type range until depletion approached 50%. As expected, as embryos approached these inflection points, we observed a peak in phenotypic variance. In the case of PAR-2 and PAR-6 (Figure 3F, 3G), asymmetry then rapidly declined as depletion was extended beyond 50%. By contrast, depletion of PAR-1 beyond 50% yielded less striking changes (Figure 3H). A shoulder is still evident as dosage levels cross the ∼50% level, but asymmetry declined only weakly thereafter (see Discussion).

Overall, we find that division asymmetry is robust to variation in PAR protein dosage, with asymmetric division phenotypes remaining at or near wild type for depletion of individual PAR proteins by up to ∼ 50%.

### Overall polarity is robust to changes in PAR dosage despite variation in local PAR concentrations

Ultimately, to achieve asymmetric division the PAR proteins must provide the appropriate spatial signals to downstream pathways. We were therefore curious how PAR protein distributions responded to dosage changes. Specifically, are distributions sensitive to changes in PAR dosage and if so, how do these changes impact the pathways that specify division asymmetry?

To this end, we quantified the distribution of PAR-2 and PAR-6 in embryos subject to progressive depletion of one or the other protein by RNAi and quantified changes in local membrane concentration and the PAR asymmetry index (ASI), in this case derived from the signal weighted contributions of *both* proteins (see Methods for ASI calculation, Figure 4A-D). We focused on embryos at nuclear envelope breakdown, when the effects of polarity cues are reduced and polarity is actively maintained by cross-talk between aPAR and pPAR proteins (Cuenca et al., 2003; Gross et al., 2019).

**Figure 4.**
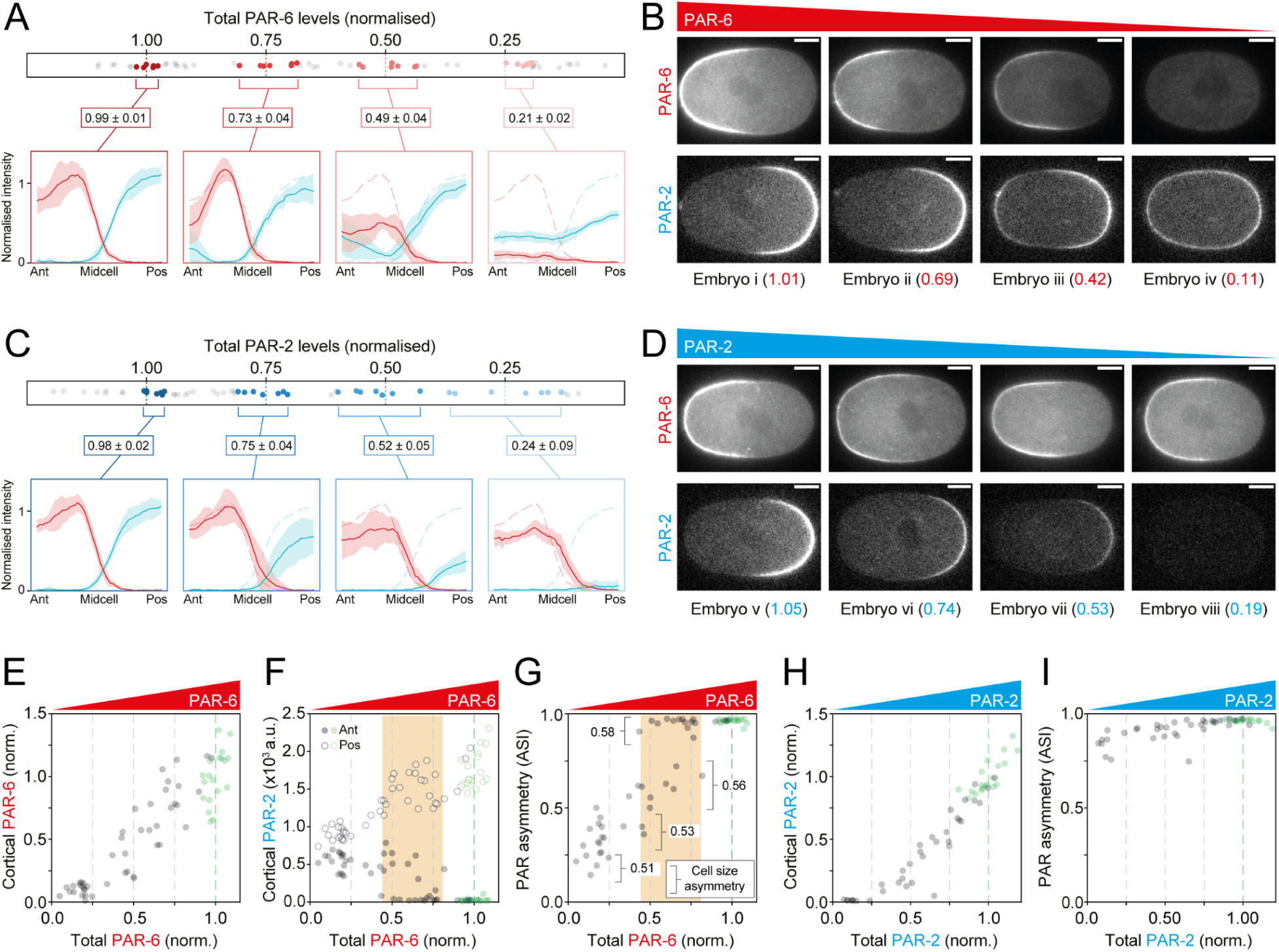
Polarity is robust despite sensitivity of the PAR network to dosage changes. **(A-D)** Evolution of PAR-2 and PAR-6 profiles as a function of the dosage of PAR-6 (A,B) or PAR-2 (C,D). Embryos expressing PAR-6::mNG and mCh::PAR-2 (NWG0268) were subject to progressive depletion of PAR-6 or PAR-2 by RNAi and dosage measured relative to mean control levels. To illustrate changes in concentration profiles, seven embryos closest to the indicated dosage levels (1.0, 0.75, 0.5, 0.25) were selected, membrane concentration profiles extracted and averaged. Mean ± SD Shown. Dashed lines in 0.75, 0.5 and 0.25 dosage profiles are the mean profiles for dosage = 1.0 for comparison. Sample embryos from the distributions in (A, C) shown in (B, D), with the dosage of the relevant PAR protein indicated. **(E)** Membrane concentrations of PAR-6 decline linearly with total PAR-6 dosage. **(F)** Reductions in PAR-6 allow invasion of PAR-2 at the anterior pole. **(G)** The relationship between PAR asymmetry (ASI) and PAR-6 dosage is bimodal. For dosage >0.75 all embryos exhibit normal asymmetry. As PAR-6 levels drop below 0.75, there is a population of embryos that retain normal asymmetry (ASI > 0.9), but a second population appears in which PAR asymmetry is reduced (ASI < 0.75) and varies linearly with PAR-6 dosage. Below dosage ∼ 0.5, the population exhibiting normal asymmetry disappears. Note numbers indicate mean division size asymmetry for embryos within the given ASI ranges, which are consistent with reduced ASI in *par-6(RNAi)* embryos affecting division asymmetry. Range of PAR-6 dosages exhibiting bimodal phenotypes in (F-G) highlighted in orange. **(H)** Membrane concentrations of PAR-2 decline linearly with total PAR-2 dosage. **(I)** Overall PAR asymmetry (ASI) is only weakly affected by PAR-2 reductions due to the stability of aPAR domains. Scale bars, 10 µm.

We found that progressive reduction of either PAR-6 or PAR-2 dosage was accompanied by a steady reduction in membrane concentration within their respective domains (Figure 4A-E, 4H, S2A-B). We obtained similar results for PAR-1, PAR-3 and PKC-3, with heterozygous embryos exhibiting roughly 50% reductions in concentrations at the membrane (Figure S2C-E). Thus, there does not appear to be any mechanism to stabilize membrane concentrations in the face of changing dosage such as occurs in wave pinning-like models for polarity where changes in boundary position can buffer the effects of dosage changes on membrane concentrations (Mori et al., 2008).

The progressive reduction of PAR-6 and PAR-2, in turn, impacted the resulting patterns of PAR protein localization, particularly as relevant dosage was reduced below 50% (Figure 4A-D). As a given PAR protein is depleted, the opposing PAR species progressively invades its domain, though the nature of this invasion differs between the two classes of PAR proteins. When PAR-2 is depleted, the anterior PAR-6 domain expands into the posterior. This expansion is accompanied by overall reductions in PAR-6 membrane concentration as PAR-6 must occupy a larger area, but PAR-6 does not become fully uniform. Thus embryos remain grossly polarized (ASI > 0.75) across the full range of PAR-2 concentrations (Figure 4C, 4D, 4I). This is consistent with reports of additional stabilizing features that limit posterior spread of aPARs, including other pPARs that act during the polarity maintenance phase, PAR-dependent cortical flows that stabilize PAR domain boundaries, and positive feedback among aPAR proteins (Kumfer et al., 2010; Lang et al., 2023; Sailer et al., 2015). Despite the relative stability of the PAR-6 domain, the PAR-2 domain gradually shrinks, becoming undetectable as PAR-2 dosage is reduced below ∼25% (Figure 4C, 4D, 4H).

By contrast, as PAR-6 was depleted, we observed the formation of a second PAR-2 domain in the anterior, presumably due to the reduced ability of aPARs to fully exclude PAR-2 from the anterior membrane (Figure 4A, 4B, 4F). Anterior domains are thought to reflect the response of PAR-2 to secondary cues that are normally suppressed in wild-type embryos (Klinkert et al., 2019; Reich et al., 2019). The presence of a second competing PAR-2 domain became increasingly frequent as depletion approached 50% (Figure 4A, 4B, 4F). For intermediate levels of depletion of between 25-75%, the behavior of the network was bimodal (Orange highlights, Figure 4F, 4G, S2F): For similar levels of depletion, one population of embryos maintained effectively wild-type levels of PAR asymmetry (ASI > 0.9). The second exhibited reduced asymmetry (ASI < 0.75) which correlated with PAR-6 concentrations, suggesting a direct relationship between aPAR activity and the relative amounts of PAR-2 at the two poles in this regime. This switch between populations seems consistent with a minimum threshold level of anterior aPAR activity required to reliably exclude PAR-2. Such a threshold would help ensure that most embryos maintain normal levels of PAR asymmetry (ASI > 0.9) so long as PAR-6 dosage remains above 50% (Figure 4G). The relationship between PAR asymmetry and division asymmetry was also highly nonlinear for PAR-6 depleted embryos with the division asymmetry of zygotes collapsing abruptly as PAR asymmetry dropped below ASI ∼ 0.5 (Figure S2G,S2H).

Taken together, our data suggest that overall PAR asymmetry (at least as reflected in the ASI) is generally robust to relative depletion of PAR proteins by up to ∼50%. Such robustness of asymmetry is likely at least part of the answer as to why asymmetric division phenotypes are robust to dosage changes. At the same time, dosage reductions are associated with progressive changes in other quantitative features of PAR protein localization, such as concentration profiles, peak membrane concentrations, domain boundaries, and levels of PAR proteins in the ‘wrong’ domain. Thus, the PAR network is clearly sensitive to alterations in PAR protein dosage, even within the regime in which overall polarity is maintained and division asymmetry is normal. These observations therefore raise questions regarding the critical signals provided by the PAR network to downstream pathways, and how they are interpreted to ensure robust outcomes in the face of quantitative changes in PAR outputs, a topic we address in the next section.

### Asymmetric division pathways canalize variation in cortical PAR input signals

Current models for PAR-dependent asymmetric division invoke local concentrations of PAR proteins as the key signals regulating division asymmetry pathways (Galli et al., 2011; Griffin et al., 2011). Yet the phenotypic endpoints of asymmetric division remain wild-type despite significant changes in local PAR concentration. How can we square these observations? Despite substantial insight into the molecular mechanisms that underlie asymmetric division, we have little quantitative data on how polarity is interpreted by downstream pathways. In particular, to what degree do downstream pathways ‘see’ changes in local PAR concentrations? Or are they only sensitive to large scale changes in polarity? We therefore sought to characterize intermediate functional readouts of the pathways underlying size and fate asymmetry.

Size asymmetry of P0 daughters arises from posterior spindle displacement during late metaphase/anaphase of P0 mitosis (Rose and Gonczy, 2014). Posterior spindle displacement shifts the division plane towards the posterior by a characteristic distance, thereby creating a smaller P1 and larger AB cell. Displacement is induced by PKC-3-dependent asymmetries in the number and/or activity of cortical force generators consisting of dynein, LIN-5(NuMa), GPR-1/2(LGN), and Gαi, which exert a pulling force on astral microtubules that reach the cortex (Colombo et al., 2003; Galli et al., 2011; Gotta et al., 2003; Gotta and Ahringer, 2001; Grill et al., 2003, 2001; Lorson et al., 2000; Srinivasan et al., 2003).

To probe sensitivity of the spindle positioning pathway, we used embryos heterozygous for PAR-2 or PAR-6 to achieve intermediate dosage reductions in a regime that does not impact size asymmetry. We then quantified key readouts of the spindle positioning pathway, including spindle elongation, spindle displacement along the A-P axis, and transverse spindle oscillations. Transverse oscillations arise during spindle elongation and are thought to depend on a critical threshold pulling force, and thus serve as a sensitive readout of changes in the forces applied to spindles (Pecreaux et al., 2006).

We found that the rate and degree of spindle elongation as well as final spindle position were nearly identical to wild-type in both *par-2* and *par-6* heterozygotes (Figure 5A-B). The only visible difference was a significant reduction in the magnitude of posterior spindle oscillations in *par-2* heterozygotes (Figure 5C-D). The effect on spindle oscillations implies that the forces acting on the spindle may be sensitive to modest reductions in PAR-2 dosage, consistent with concentration-dependent regulation of components of the spindle positioning apparatus. At the same time, because the 30-40% reductions in PAR concentrations in *par* heterozygotes did not impact the speed of spindle elongation or accuracy of positioning along the A-P axis, there must be substantial nonlinearity in the readout of local PAR concentrations by spindle positioning pathways. This could occur in the readout of PAR concentrations by regulators of cortical force generators, effectively maintaining the asymmetry in pulling forces at wild-type levels, or alternatively, spindle elongation and positioning pathways could themselves be buffered against changes in the asymmetry of pulling forces (see Discussion).

**Figure 5.**
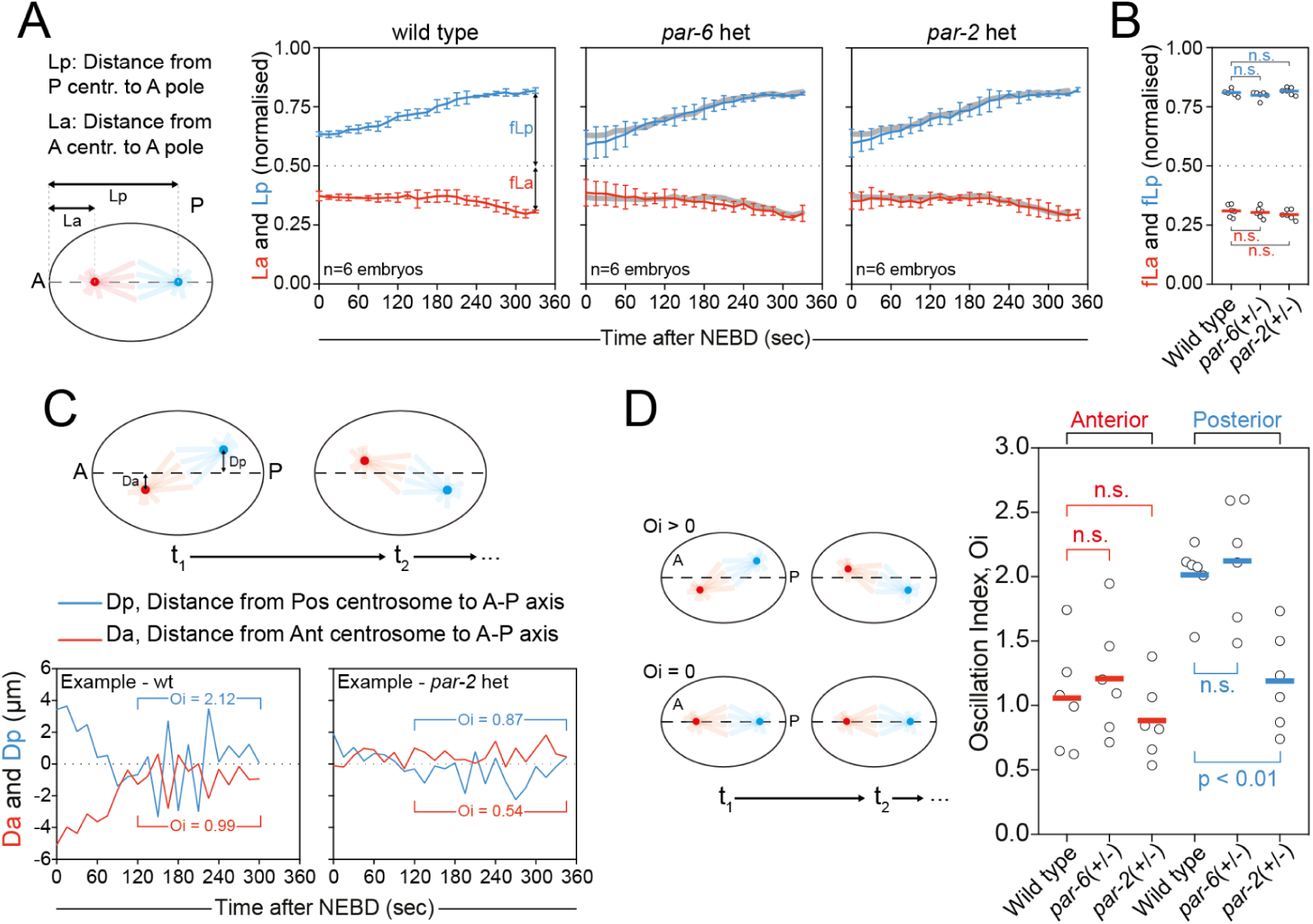
Spindle positioning is highly robust to PAR dosage changes. **(A)** Evolution of anterior and posterior centrosome position from NEBD through telophase. Mean behavior for wild-type embryos shown as gray lines in *par-2(+/-)* and *par-6(+/-)* heterozygote plots. Note nearly identical behavior in wild-type and embryos heterozygous for mutations in *par-2* or *par-6.* Mean ± SD shown. **(B)** Comparison of final centrosome positions (fLa, fLp) taken at telophase from experiments in (A), defined as the time when the cleavage furrow was 50% ingressed and centrosomes exhibited no further outward motion. Individual data points and mean indicated. **(C)** Schematic for quantifying spindle oscillations along with two example traces for embryos from wild-type or heterozygous *par-2* animals. Oscillation index (Oi) was defined as the standard deviation of Da or Dp during the period of prometaphase to telophase. **(D)** Heterozygous *par-2* embryos exhibit reduced posterior Oi relative to wild-type and *par-6* heterozygotes. Anterior Oi is similar across all three conditions. Individual data points and mean indicated. Samples in (D) were compared statistically using an unpaired t-test.

We next turned to the fate segregation pathway, which relies on the establishment of an anterior-to-posterior gradient of MEX-5/6 by a complementary gradient of PAR-1 kinase activity, which is set up downstream of cortical polarity and locally modulates the diffusivity of MEX proteins (Daniels et al., 2010; Griffin et al., 2011; Tenlen et al., 2008). The resulting MEX gradient induces asymmetric segregation of fate determinants, including various germline markers such as PIE-1 and P granules as well as the cell cycle regulators that are responsible for division asynchrony (Figure 6A)(Brauchle et al., 2003; Budirahardja and Gönczy, 2008; Han et al., 2018; Kipreos and van den Heuvel, 2019; Michael, 2016; Nishi et al., 2008; Rivers et al., 2008; Schubert et al., 2000; Wu et al., 2015).

**Figure 6.**
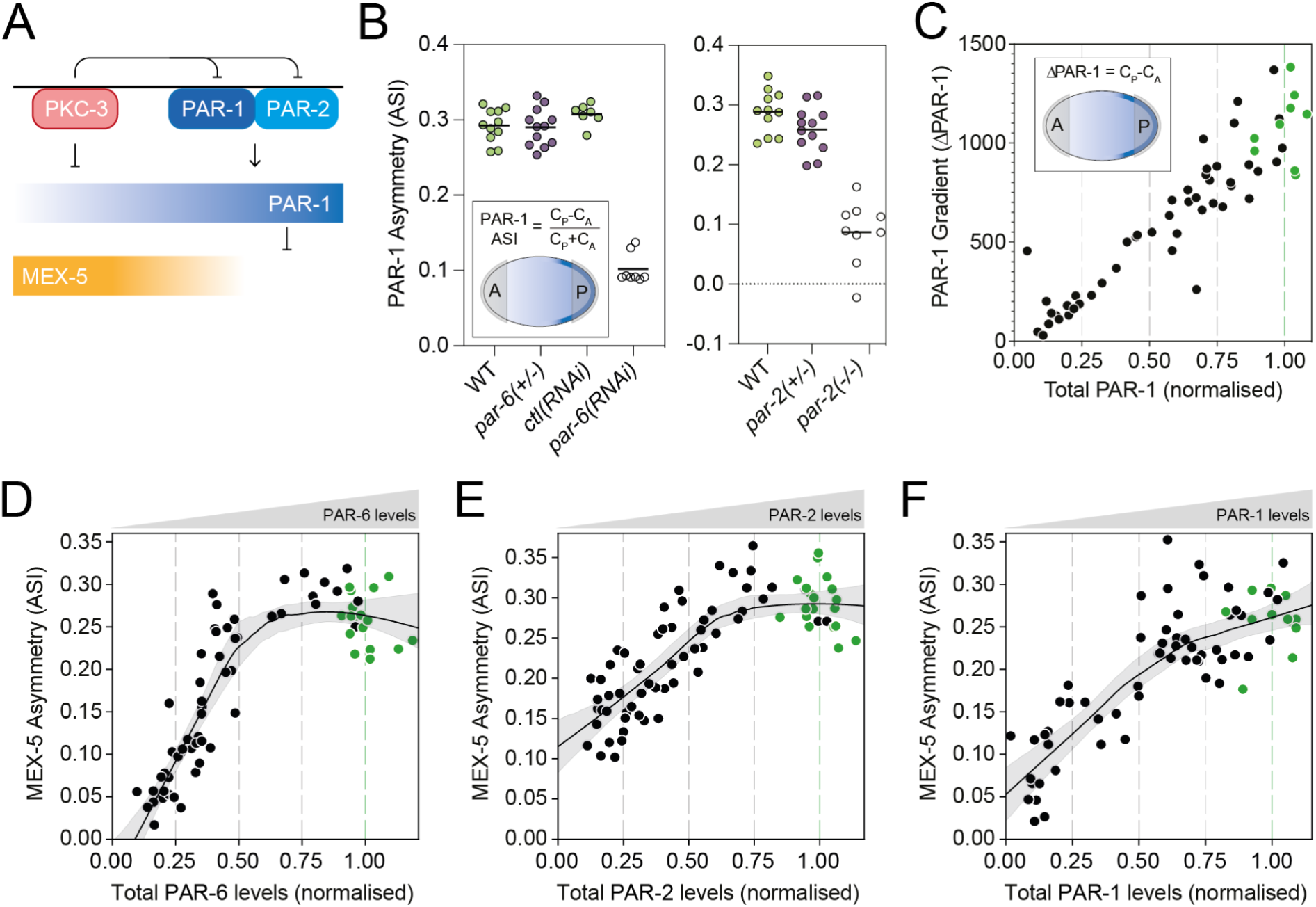
Cytoplasmic asymmetry is robust to perturbations of PAR protein concentrations. **(A)** Fate asymmetry is specified by a cytoplasmic gradient of MEX-5 that is downstream of PAR polarity and induces asymmetric inheritance of fate determinants between AB and P1 cells. Cell cycle asynchrony is a commonly used proxy of fate asymmetry. Note that the mechanistic relationship and contributions of cortical vs. cytoplasmic PAR-1 asymmetry are not well understood. **(B)** PAR-1 gradient asymmetry (ASI) is robust in *par-2(+/-)* and *par-6(+/-)* heterozygotes. Genotypes indicated. *par-6(RNAi)* and *par-2(-/-)* homozygous mutants shown for comparison. **(C)** The absolute magnitude of the PAR-1 gradient (ΔPAR-1) declines near linearly with PAR-1 dosage. Magnitude of PAR-1 concentration difference between anterior and posterior (C_P_-C_A_) shown as a function of PAR-1 dosage. **(D-F)** MEX-5 asymmetry responds nonlinearly to depletion of PAR proteins. MEX-5 asymmetry (ASI) as a function of dosage of PAR-6(D), PAR-2(E), and PAR-1(F). Fit lines in (D-F) indicate Lowess fit with 95% confidence interval determined by bootstrapping. In (C-F), wild-type data points are indicated in green.

Similar to our analysis of spindle positioning, we sought to measure proximal features of the fate segregation pathway, including the PAR-1 and MEX-5 gradients, summing PAR-1 signal from both membrane and cytoplasm for this purpose. Whereas loss of PAR-6 or PAR-2 substantially reduced or eliminated PAR-1 asymmetry, asymmetry was normal in both *par-2* and *par-6* heterozygotes (Figure 6B). Consistent with the robustness of PAR-1 asymmetry in *par-2* and *par-6* heterozygotes, when we subjected embryos to progressive depletion of either protein, the MEX gradient was unchanged until dosage declined below 50% (Figure 6D-E). We conclude that an intrinsic stability of PAR-1 asymmetry to changes in cortical PAR concentrations explains a substantial fraction of the robustness of fate asymmetry to intermediate changes in PAR dosage.

However, while this intrinsic stability of PAR-1 asymmetry can explain the robustness of fate specification with respect to PAR-2 and PAR-6 dosage, it does not explain why fate is also relatively robust to depletion of PAR-1 itself as changes in PAR-1 concentration ought to directly impact the local kinase activity available to polarize MEX-5. We therefore scored the PAR-1 and MEX-5 gradients as a function of PAR-1 dosage. As expected, the magnitude of the PAR-1 concentration difference across the zygote declined as a linear function of PAR-1 dosage (Figure 6C). By contrast, the MEX-5 gradient initially showed relatively modest changes in response to PAR-1 depletion, only decaying more strongly as depletion exceeded 50% (Figure 6F). Thus, together with the data above, our results indicate that both PAR-1 and MEX-5 exhibit significant robustness in their ability to maintain asymmetric gradients with respect to perturbations of upstream signals (i.e. cortical PAR proteins).

We therefore conclude that within the ‘robust regime’ of PAR dosage changes, both spindle and fate asymmetry pathways possess mechanisms to canalize phenotypic outputs in response to variation in local cortical PAR concentrations. Consequently, the signals provided by the cortical PAR network and the downstream asymmetric division pathways are quantitatively decoupled, helping to ensure that the precision of size and fate asymmetries is robust to variation in PAR dosage.

### PAR dosage alters sensitivity to symmetry-breaking cues

Our data so far suggest that robust division asymmetry emerges from two features: First the stability of overall PAR asymmetry against variations in PAR dosage, and second, readout mechanisms that quantitatively decouple dosage-dependent variation in PAR concentrations at the membrane from pathway outputs. We next turned our attention to symmetry-breaking to determine whether such quantitative decoupling may be a general feature of the processes leading to asymmetric division.

Symmetry-breaking is triggered by several semi-redundant symmetry-breaking cues. The dominant cue is cortical actomyosin flow, which is induced by the centrosome and advects aPAR proteins into the nascent anterior allowing pPAR proteins to invade the posterior membrane (Goehring et al., 2011; Munro et al., 2004). Coincidentally, a second, flow-independent centrosomal cue locally promotes pPAR loading onto the posterior pole through a pathway that involves protection of PAR-2 by centrosomal microtubules (Motegi et al., 2011; Zonies et al., 2010). Finally, a number of other cues have also been proposed that may enhance the efficiency of symmetry-breaking, including curvature and hydrogen-peroxide produced by centrosome-associated mitochondria (De Henau et al., 2020; Klinkert et al., 2019). While the existence of multiple cues may help explain why symmetry-breaking is robust to loss of a given cue, how the strength of symmetry-breaking cues is related to the resulting polarity of the PAR network is less clear.

Theoretical models suggest that the self-organizing properties of the PAR network allow it to convert potentially variable symmetry-breaking cues into stable and reproducible polarity outputs (Goehring et al., 2011; Gross et al., 2019)(Schematic in Figure 7A). Specifically, once sufficient asymmetry is imparted to the PAR network to break symmetry, feedback within the PAR network takes over to drive the system to a parameter-defined polarized steady state. At the same time, the magnitude of asymmetry induced by so-called “guiding cues” in wild-type embryos is thought to be well in excess of that required to break symmetry (Gross et al., 2019). Such strong guiding cues likely insulate the embryo against variance in PAR feedback that otherwise could impact the reliability of symmetry-breaking. Such models therefore predict that we should observe nonlinear relationships between the strength of symmetry-breaking cues and the resulting polarity of the zygote, similar to what we observed between PAR polarity and division asymmetry. Specifically, we predict that polarity outputs in otherwise wild-type embryos should be robust to changes in cue strength and, conversely, that reductions in cue strength should render polarity outputs sensitive to PAR dosage.

**Figure 7.**
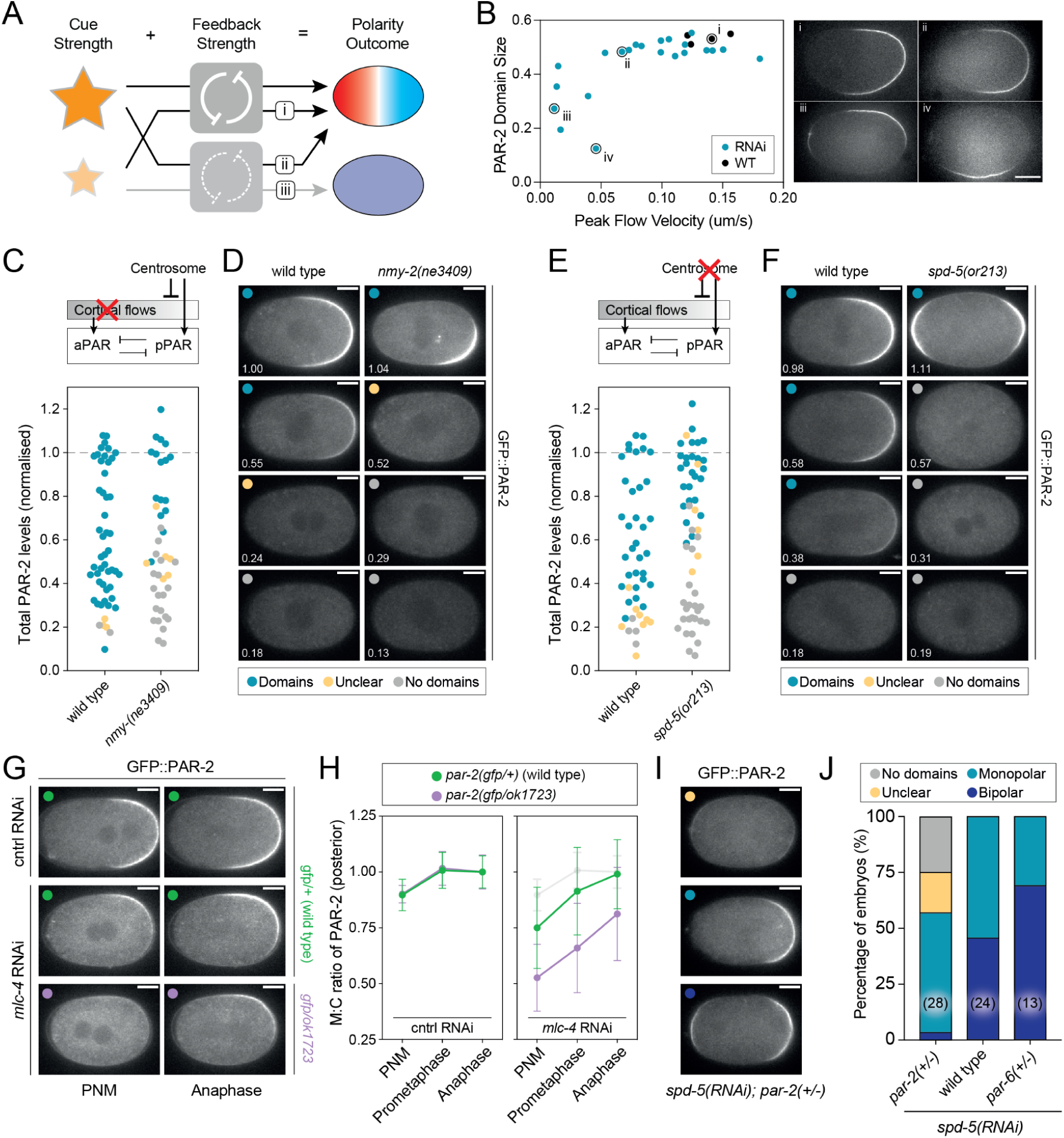
PAR dosage reduction renders polarity sensitive to defects in polarity cues. **(A)** Scheme for how reduced PAR dosage could sensitize embryos to compromised symmetry-breaking cues. At full strength, PAR feedback is sufficient to amplify signals provided by a weakened cue and thereby rescue normal polarity (i). Conversely,sufficiently strong cues can compensate for reduced PAR feedback to rescue polarity establishment in embryos partially depleted of PAR proteins (ii). However, in the presence of reduced PAR feedback, polarity becomes sensitive to cue strength (iii). **(B)** Above a threshold cortical flow velocity, PAR-2 domain size is nearly constant. Only upon progressive reduction in peak cortical flow velocity below a threshold value do PAR-2 domains undergo an abrupt shift to being variable sized and mispositioned, consistent with a shift to a flow-independent symmetry-breaking regime. Cortical flow velocities were reduced by *mlc-4(RNAi)*. Individual embryos are indicated and images of select examples shown at right. **(C-D)** Inhibition of cortical flow sensitizes symmetry-breaking to reduced PAR-2 dosage. GFP::PAR-2 dosage was progressively reduced by RNAi in wild-type and temperature-sensitive *nmy-2(ne3409ts)* embryos at the restrictive temperature (25°C) and embryos imaged just prior to NEBD. PAR-2 dosage was measured, embryos scored for the presence of GFP::PAR-2 domains, and the results plotted in (C). Example embryos at different PAR-2 dosages and exhibiting different phenotypes shown in (D). **(E-F)** Disruption of the centrosome cue sensitizes symmetry-breaking to PAR-2 dosage. Performed as in (C-D), but using the temperature-sensitive allele *spd-5(or213).* Embryos were scored as exhibiting domains if they exhibited clearly defined PAR-2 domains. Note that *spd-5(or213)* embryos often exhibit bipolar PAR-2 domains, which were scored as having domains for the purposes of this assay (see I-J). **(G-H)** Heterozygosity for *par-2(ok1723)* substantially delays symmetry-breaking in *mlc-4(RNAi)* embryos with reduced flows. Images (G) and quantification of membrane:cytoplasm ratio (H) at pronuclear meeting (PNM), prometaphase, and anaphase for *par-2(gfp/+)* or *par-2(gfp/ok1723)* embryos subject to either control or *mlc-4(RNAi)*. Control data shown in light gray in mlc-4(RNAi) panel for comparison. Note *par-2(gfp/+)* embryos were used as controls so that quantification would not be affected by differing levels of GFP signal. 3**(I-J)** Heterozygosity for *par-2(ok1723)* compromises symmetry-breaking in *spd-5(RNAi)* embryos with defective centrosomes. *spd-5(RNAi)* embryos with the indicated *par* genotypes were scored for the presence or absence of clearly defined GFP::PAR-2 domains and whether one (monopolar) or two (bipolar) domains were present (J). Example images of scored phenotypes in (I). Scale bars, 10µm.

To directly test these hypotheses, we first measured PAR-2 domain size as a function of cortical flow velocity. Flow velocity was tuned through RNAi-mediated depletion of the myosin regulatory light chain, MLC-4. We found that domain size at NEBD was remarkably constant until cortical flow velocities were reduced by over 50% (below 0.05 µm/s). However, beyond this point domains became highly variable in size and position (Figure 7B). This increased variability likely reflects a transition to flow-independent symmetry-breaking pathways, which have previously been associated with delays in domain formation, slow domain expansion, and failure to align polarity domains with the long axis (Goehring et al., 2011; Goldstein and Hird, 1996; Gross et al., 2019; Motegi et al., 2011; Zonies et al., 2010). Consistent with this interpretation, the observed threshold velocity of 0.05 um/s roughly corresponds to the minimal cortical flow velocity required for symmetry-breaking when flow-independent cues are compromised(0.062 µm/s) (Gross et al., 2019). Thus, while embryos require a minimal flow velocity to enter a flow-dependent polarization regime, once this is achieved, the degree of PAR polarity - here defined by PAR-2 domain size - exhibits minimal variability and is largely decoupled from the strength of the symmetry-breaking cue.

We next asked whether reduction in the magnitude of guiding cues rendered embryos sensitive to dosage variation by assessing polarity as a function of PAR-2 dosage under two distinct symmetry-breaking regimes: a flow-defective regime (*nmy-2(ne3409)* or *mlc-4(RNAi)*) in which flows are absent and polarity is thought to rely on centrosomal microtubules (Gross et al., 2019; Motegi et al., 2011; Zonies et al., 2010) and a centrosome-defective regime (*spd-5(or213)* or *spd-5(RNAi)*) in which the posterior centrosomal cue is compromised (Hamill et al., 2002; Kapoor and Kotak, 2019; Klinkert et al., 2019; Liu et al., 2004; O’Connell et al., 2000; Sonneville and Gönczy, 2004; Tsai and Ahringer, 2007; Zhao et al., 2019). We found that control embryos with intact symmetry-breaking cues reliably formed PAR-2 domains despite reduction of PAR-2 dosage by up to ∼75% relative to wild-type levels. By contrast, in both flow- and centrosome-defective conditions, PAR-2 domain formation failed when depletion exceeded 30-40%, marking a clear shift in the threshold level of PAR-2 required for efficient symmetry-breaking (Figure 7C-F).

Corroborating our PAR-2 rundown experiments, PAR-2 domains were weaker and delayed in *par-2* heterozygous embryos subject to *mlc-4(RNAi)* compared to wild-type controls (Figure 7G, 7H). As we showed (see Figure 2C), *par-2* heterozygotes contain PAR-2 amounts that are roughly 60-70% of those found in wild-type embryos, which corresponds roughly to the point in the PAR-2 rundowns at which we began to observe polarization defects in cue-compromised embryos (Figure 7C).

Finally, to explicitly test the effects of both increased and decreased aPAR:pPAR ratios in cue-compromised conditions, we compared polarity outcomes in wild-type, heterozygous *par-2*, and heterozygous *par-6* embryos depleted of SPD-5. We found that outcomes strongly depended on dosage (Figure 7I, 7J): in *spd-5(RNAi)* embryos with normal PAR dosage, all embryos exhibited clear PAR-2 domains, with roughly a 50:50 mix between monopolar embryos with a single PAR-2 domain, and bipolar embryos with PAR-2 domains at both anterior and posterior poles. As noted above, bipolar embryos are thought to arise from the loss of a single dominant centrosomal cue at one pole, which results in the embryo responding inappropriately to weak cues at the anterior pole that are normally ignored (Klinkert et al., 2019; Reich et al., 2019). By contrast, in *spd-5(RNAi)* embryos heterozygous for *par-2*, roughly half of embryos failed to exhibit clear PAR-2 domains and among embryos with a PAR-2 domain, bipolarity was rare. Thus, as in our RNAi experiments, reduced PAR-2 dosage compromises the ability of embryos to respond to sub-optimal symmetry-breaking cues, including weak anterior cues that can lead to bipolarity in *spd-5* embryos. Strikingly, *par-6* heterozygotes exhibited the opposite trend - all embryos exhibited clear PAR-2 domains, but the majority of embryos were bipolar. Thus, our data highlight how strong guiding cues effectively mask an underlying sensitivity of the symmetry-breaking process to PAR dosage.

Overall, consistent with predictions in (Gross et al., 2019), the self-organizing features of the PAR network effectively insulates the zygote from substantial variations in the strength of symmetry-breaking cues, ensuring robust polarity outputs in the face of perturbations in polarizing cues. However, as we have shown here, this ability to canalize variation in cue strength requires normal activity of the PAR network and consequently, in embryos with compromised cues, polarity outcomes become highly sensitive to PAR dosage (Figure 7A). This last observation likely explains strain-dependent differences in polarity phenotypes in cue-compromised embryos when using different tagged alleles or ectopic transgenes (Kapoor and Kotak, 2019; Klinkert et al., 2019; Reich et al., 2019; Tsai and Ahringer, 2007; Zhao et al., 2019; Zonies et al., 2010), and could account for the acute sensitivity of polarity phenotypes in cue-compromised conditions to weak hypomorphic mutations that affect PAR-2 function (Calvi et al., 2022; Motegi et al., 2011).

## Discussion

Our data argue that neither dosage compensation nor compensatory regulation of dosage between network components are sufficient to explain robustness of asymmetric division. Rather, we find that division asymmetry remains substantially normal in the face of ∼50% reductions in protein amounts. Surprisingly, this near wild-type division asymmetry can occur despite the fact that partial PAR depletion results in changes in both the local concentration and distribution of PAR proteins, and in the strength of the feedback circuits required to promote polarization.

It was somewhat curious that both PAR-2 and PAR-6 were expressed at modestly higher levels than expected levels in heterozygotes, which in principle would help ensure that levels in heterozygotes remain above the level where one starts to see defects in division asymmetry. However, embryos heterozygous for *par-1*, *par-3*, and *pkc-3* did not exhibit such increases (Figure 1D), yet still showed similar near wild-type asymmetric division phenotypes, suggesting that partial upregulation is neither a general adaptation of *par* genes nor generally required for the stability of phenotypes in *par* heterozygotes.

Instead, our data reveal nonlinear relationships between signal inputs and output responses as a core feature of the asymmetric division pathway. These nonlinearities effectively canalize variation allowing embryos to achieve near wild-type levels of accuracy despite variability in the function or activity of a given module. Specifically, a wild-type PAR network can generate normal polarity in the face of weakened symmetry-breaking cues, while spindle positioning and fate segregation pathways can achieve reproducible division asymmetry despite broad variation in the quantitative parameters of PAR polarity such as domain size and local PAR concentrations.

Through such nonlinear input-output maps, ‘modules’ for symmetry-breaking, PAR polarity, and spindle/fate asymmetry are quantitatively decoupled, minimizing the effects of genetic, environmental, or stochastic variation at each step. We hypothesize that the ability of each module to ‘correct’ for defects arising in earlier modules has the effect of preventing error from accumulating as the embryo proceeds along this developmental trajectory leading to stability of phenotypic outcomes.

An important consequence of such a regime is that quantitative measures of division asymmetry such as size and fate are not directly set by the strength of symmetry-breaking cues or the magnitude of specific features of the PAR pattern such as domain size or PAR boundary position. In this sense, the PAR pattern does not encode concentration-dependent positional information and thus is dissimilar to models of position specification by gradients (Briscoe and Small, 2015; Hubatsch and Goehring, 2020). Instead, we observe a more steplike collapse of division asymmetry in response to dosage perturbations, which seems most consistent with a requirement for a minimal threshold magnitude of polarity to define the polarity axis.

One common explanation for robustness to dosage changes is that enzymes are simply present in excess. Genetic dominance describes the observation that the vast majority of wild-type alleles are ‘dominant’ over loss-of-function alleles (Deutschbauer et al., 2005; Fisher, 1928). To explain the origin of dominance, it has been postulated that insensitivity to 2-fold reductions in enzyme concentration would likely be selected for as a safety feature unless a pathway is sensitive to overexpression (Haldane, 1939; Kacser and Burns, 1981; Wright, 1934). Consistent with this idea, work in *S. cerevisiae* demonstrated that sensitivity to 2-fold reductions in dosage (i.e. haploinsufficiency) results from the relatively rare cases in which both over- and under-expression impose substantial fitness costs (Morrill and Amon, 2019). A priori, the requirement for balance between opposing PAR species could have led one to expect sensitivity to dosage. Indeed, if one looks only at the precision of the PAR pattern, such as local PAR concentrations or PAR domain size, the PAR network is sensitive to dosage changes. Yet, when we ask how polarity is interpreted, it is apparent that downstream processes are relatively insensitive to these quantitative variations in polarity signals. Thus, the asymmetric division pathway in the *C. elegans* zygote appears similar to many signaling networks in having evolved nonlinear or threshold-like responses to ensure robust phenotypic outcomes (e.g. asymmetric division) in the face of input signal variation (e.g. cue strength or local PAR concentrations).

A key question that remains is how nonlinear input:output mapping is encoded. The self-organizing properties of the PAR network provide one answer (Arata et al., 2016; Bland et al., 2023; Goehring et al., 2011; Gross et al., 2019; Lang et al., 2023; Sailer et al., 2015). Self-organization relies on bistable reaction kinetics, which endow the system with switch-like behaviors and threshold-like responses that are commonly associated with robustness. Although the extent of feedback within the PAR network remains an open question (Goehring, 2014; Lang and Munro, 2017; Motegi and Seydoux, 2013), these basic properties provide a strong driving force towards a stable polarized steady-state (this work and (Gross et al., 2019)). Biochemical driving forces are known to be further complemented by boundary stabilizing effects of patterned actomyosin flows induced by the PAR proteins themselves (Sailer et al., 2015). As we show, these features allow the system to cope with substantial perturbations, such as the strength of the symmetry-breaking cues, but also to maintain the stability of overall polarity with respect to PAR dosage.

By contrast, the design features that provide for robustness in the interpretation of variable PAR polarity signals by downstream pathways are less clear. As the precise mechanisms by which PAR signals are transduced to downstream spatial pathways remain enigmatic, a full exploration of the features in these pathways that give rise to nonlinear readouts is beyond the scope of this work. However, it is striking how little these processes are affected by real and substantial alterations in PAR concentrations given that spindle pulling forces and cytoplasmic gradients of fate determinants are thought to be directly regulated by the PAR kinases PKC-3 and PAR-1.

In the case of spindle positioning, key regulators of cortical force generators such as LET-99 and LIN-5 are thought to be targets of polarity kinases (Galli et al., 2011; Grill et al., 2001; Wu and Rose, 2007). Yet we find that spindle position is largely robust to 50% changes in the concentration of polarity kinases. In fact, it is curious that spindle position appears less sensitive to the kinase PAR-1 than PAR-2, whose primary role has been linked to stabilizing PAR-1 at the posterior membrane, perhaps suggesting a more direct role for PAR-2 in regulating spindle forces. Such a result would be consistent with quantitative trait mapping that linked *par-2* with spindle size (Farhadifar et al., 2020).

Previous work has shown that spindle positioning is robust to changes in absolute magnitude of pulling force, even under conditions that alter the rate of spindle elongation and/or posterior displacement (Pecreaux et al., 2006; Redemann et al., 2011), suggesting the existence of positional control mechanisms (Bouvrais et al., 2018; Farhadifar et al., 2020; McCarthy Campbell et al., 2009). Here we show that mechanisms must also exist to render positioning processes insensitive to quantitative alterations in the relative PAR concentrations along the A-P axis, given that, aside from the reduction in spindle oscillations in *par-2* heterozygotes, the pattern of spindle elongation and posterior spindle pole displacement were quantitatively unchanged in *par* heterozygotes.

Given that the PAR-1 gradient is largely unaffected by heterozygosity in either *par-6* or *par-2*, and MEX-5 is in turn resilient to partial depletion of its immediate upstream regulator PAR-1, cytoplasmic gradients are also unlikely to simply reflect readout of local concentrations of the relevant polarity kinases. Given our observations that PAR-2 invasion of the anterior was associated with depletion of aPARs below a critical level, it is tempting to speculate that threshold-like responses to local PAR kinase activity may be a general principle to enable robust asymmetric regulation of downstream targets, including regulators of cortical force generation and cytoplasmic asymmetries. How these threshold behaviors are encoded will be a key area for future investigation.

In conclusion, through establishing quantitative perturbation-phenotype maps for the first embryonic cell division of *C. elegans*, we have revealed how nonlinear readouts of spatial information drive robust developmental outcomes within an asymmetric division program.

## Methods

### C. elegans strains and maintenance

*C. elegans* strains were maintained on OP50 bacterial lawns seeded on nematode growth media (NGM) at 16°C or 20°C under standard laboratory conditions (Stiernagle, 2006). Worm strains were obtained from *Caenorhabditis* Genetics Center (CGC) and are listed in Supplemental Table S1. Note that analysis of zygotes precludes determination of animal sex.

### C. elegans husbandry and generation of heterozygous genotypes

To generate fluorescently-tagged heterozygous animals harboring one mutant allele copy, males of relevant balanced strains were crossed into hermaphrodite L4 larvae of homozygous FP-tagged strains (see Figure 2B-D, 3C-D, 3F-G, 6B, 7G-J). Similarly, to obtain heterozygous worms where the two alleles of a given gene are tagged with different FPs, males of a given homozygous background (e.g., mCherry-tagged gene) were crossed into hermaphrodite L4 larvae of a homozygous strain expressing the relevant gene labeled with a different FP (e.g., GFP) (see Figure 2D, S1, S2A-E). To study polarity in specimens expressing an FP-tagged par in contexts where a different par is present at varying dosage, larvae of either heterozygous or null genotypes were directly selected from balanced strains prior to sample preparation and imaging (see Figure 3H-I).

### RNAi culture conditions

RNAi by feeding was performed according to previously described methods (Kamath and Ahringer, 2003). Briefly, HT115(DE3) bacterial feeding clones were inoculated from LB agar plates to LB liquid cultures and grown overnight at 37°C in the presence of 50 µg/mL ampicillin (until a fairly turbid culture is obtained). To induce high dsRNA expression, bacterial cultures were then treated with 1 mM IPTG before spotting 150 µL of culture onto 60 mm NGM agar plates (supplemented with 10 µg/ml carbenicillin, 1 mM IPTG) and let incubate for 24 hr at 20°C. In general, to obtain strong/complete gene depletion (or allele-specific depletion in the case of experiments using dual-labeled alleles), L3/L4 larvae were added to RNAi feeding plates and incubated for 24-36 hr depending on gene and temperature (see Figures 2D, 2G-H, S1, S2A-E, 6B, and 7G-J). To perform protein rundowns, where dosage is depleted from wild type concentration through full depletion, L3/L4 larvae were either: placed on relevant RNAi for variable periods of time (from 4 hr to 24-36 hr incubation); or placed on plates where the relevant RNAi feeding clone was mixed with a control RNAi clone (expressing non-targeting dsRNA) to dampen the strength of protein depletion (see Figures 2E-F, 3C-H, 4, S2A-B, S2F-H, 6C-F and 7B). For the protein rundowns using temperature-sensitive strains, L3/L4 larvae were placed on RNAi for variable amounts of time (or on plates containing par RNAi mixed with control RNAi, as detailed above) at 16°C, and then placed on OP50 and shifted to 25°C for 75 min (see Figures 7C-D) or 45 min (see Figures 7E-F) prior to imaging.

### C. elegans dissection and mounting for microscopy

For most experiments (namely, for Figures 2B-F, S1, 3, 4, S2, 5, 6D-F, 7B), early embryos were dissected from gravid hermaphrodites in 5-6 µL of M9 buffer (22 mM KH_2_PO_4_, 42 mM NaHPO_4_, 86 mM NaCl and 1 mM MgSO_4_) on a coverslip and mounted under 2% M9 agarose pads (Zipperlen et al., 2001). In some instances (see Figures 2G-H, 6B-C, 7C-J), to minimize eggshell autofluorescence that can be prominent with agarose mounts, embryos were dissected in 8-10 µL of egg buffer (118 mM NaCl, 48 mM KCl, 2 mM CaCl_2_ 2 mM MgCl2, 25 mM HEPES, pH 7.3), and mounted with 20 µm polystyrene beads (Polysciences, Inc.) between a slide and coverslip as in (Rodriguez et al., 2017).

### Microscopy

Unless specified otherwise, midsection confocal images were captured on a Nikon TiE with a 60x/1.40 NA oil objective, further equipped with a custom X-Light V1 spinning disk system (CrestOptics, Rome, Italy) with 50 µm slits, Obis 488/561 fiber-coupled diode lasers (Coherent, Santa Clara, CA) and an Evolve Delta EMCCD camera (Photometrics, Tuscon, AZ). Imaging systems were run using Metamorph (Molecular Devices, San Jose, CA) and configured by Cairn Research (Kent, UK). Filter sets were from Chroma (Bellows Falls, VT): ZT488/561rpc, ZET405/488/561/640X, ET535/50m, ET630/75m.

To obtain confocal images of samples expressing mNG-tagged genes, imaging was performed on a Leica TCS SP8 inverted microscope (Leica Microsystems Ltd, Wetzlar, Germany), equipped with an Apo CS2 63x/1.40 NA oil objective and a HyD detection system. Imaging was managed with LAS X software (Leica Microsystems Ltd, Wetzlar, Germany), and acquisition was set at a scanning speed of 400 Hz with pinhole aperture set to 2 AU. Unlike the Nikon configuration detailed above, this microscope offered the capability of imaging samples with either 488-nm or 514-nm excitation, thus permitting the distinction between GFP and mNG specific fluorescence (as in the case of experiments shown in Figure 2D, S1D-F, S2C-E).

For experiments shown in Figure 7B, embryos were imaged on an Olympus IX71 equipped with a Yokogawa spinning disk head using a 63x/1.40 Oil UPlanSApo objective, 488, 561-nm DPSS lasers, an iXon EMCCD camera and ImageIQ (Andor Technology).

For most experiments (see Figures 2, S1, 4, S2, 6B-C, 7B-F, 7I-J), image acquisition was performed by taking still images of live embryos at NEBD, metaphase and telophase/cytokinesis (ie, early 2-cell stage). Data were acquired with both fluorescence and transmitted light configurations. In all experiments, imaging was done at 20°C, except in the case of temperature-sensitive alleles, where acquisition was done at 25°C using an objective temperature control collar (Bioptechs) (see Figures 7C-F). In some instances (see Figures 3C-H), the imaging pipeline was expanded to include analysis of the embryonic 4-cell stage. For this, timeseries of embryo development were acquired (at 15 sec interval) as the AB and P1 blastomeres underwent cell division.

To image zygotic polarization in the various dosage/cortical flow regimes shown in Figure 7G-H, embryos were imaged from pronuclear migration/early prophase through telophase/cytokinesis at 60 s intervals.

For data shown in Figures 3A-B, where unlabeled lines were used, embryos were imaged using transmitted light only (i.e., DIC). In this case, data acquisition was only done at the 2-cell (snapshot taken at ‘birth’ of AB and P1 cells) and 4-cell stage development (performing timeseries as detailed above).

To image spindle dynamics in mitotic zygotes (as detailed in Figure 5), samples were filmed from NEBD through telophase at 15 s intervals using DIC.

### Image processing prior to fluorescence quantitation

For most experiments, and in order to quantify fluorescence signal and gauge protein dosage (total and/or cortical concentrations), images of embryos were taken alongside a local background image (with no samples in the field of view), which was subtracted from the image prior to analysis. Note that this step can usually be omitted without much detriment; however, background subtraction may improve images in cases where the background signal is uneven or variable.

### Image analysis - quantitation of total and cortical fluorescence

For quantification of whole-embryo fluorescence intensities (to account for protein levels in both cytoplasm and membrane/cortex), mean pixel intensity was measured within a manually-defined region of interest (ROI) encompassing the entire cross-section of the embryo.

To measure cortical concentrations, a 50-pixel-wide (12.8 µm) line following the membrane around the embryo was computationally straightened, and a 20-pixel-wide (5.1 µm) rolling average filter was applied to the straightened image. Intensity profiles perpendicular to the membrane at each position were fit to the sum of a Gaussian component, representing membrane signal, and an error function component, representing cytoplasmic signal. Membrane concentrations at each position were calculated as the amplitude of the Gaussian component. Cortical levels in Figures 4E-I, S2 were calculated as the mean membrane concentration in the posterior-most (PAR-1, PAR-2), or anterior most (PAR-3, PAR-6, PKC-3) 33% of the embryo perimeter. This protocol is similar to previously published methods (Gross et al., 2019; Reich et al., 2019).

For quantification of fluorescence of GFP- or mNG-tagged proteins (when using 488-nm excitation and a 535/50 nm emission filter), we applied SAIBR for autofluorescence correction (Rodrigues et al., 2022). This protocol was established to circumvent the high autofluorescence emission that results from 488-nm excitation (the most commonly used wavelength when imaging green fluorophores). In particular, the method exploits the fact that autofluorescence typically has a much wider emission spectrum than GFP. The protocol involves the use of a parallel channel, with a red-shifted emission filter (namely, a 630/75 nm filter), that is used to gauge the level of autofluorescence in the sample. The inferred autofluorescence is then subtracted from the measured signal intensity in the GFP channel (which is, in essence, a sum of both fluorophore-specific and non-specific signals), thus yielding a more accurate estimate of protein concentrations. Subtraction can be done on a pixel-by-pixel basis (allowing for spatial signal correction) or on an embryo-by-embryo basis (e.g., to quantify whole-embryo fluorescence intensities, as indicated above). Note that SAIBR is also compatible when quantifying GFP fluorescence in embryos that co-express additional, spectrally-distinct fluorophores (such as mCherry), as detailed in (Rodrigues et al., 2022).

To quantify fluorescence in embryos expressing mNG-tagged PARs when using a 514-nm excitation and 550/50 nm emission filter configuration which minimizes autofluorescence, hereafter referred to as ‘mNG channel’, a mean background signal was first measured across a sample of unlabeled embryos, and this mean value was then simply subtracted from the mNG channel signal in the mNG-tagged embryos. A similar protocol was employed in the quantitation of mCherry-tagged PAR signal, but in this case utilizing a channel with 561-nm excitation and a 630/75 nm emission filter.

### Image analysis - dependence of PAR-2 polarity on cortical flows

For this experiment, embryos expressing NMY-2::GFP and mCherry::PAR-2 (TH306) were imaged on an Olympus IX71 spinning disk microscope (details above). Cortical flow velocities were extracted from cortical GFP timeseries data (2 s interval) by particle image velocimetry using the Matlab *mpiv* package using custom scripts. PAR-2 domain size was quantified from midplane images taken at NEBD (see Figure 7B).

### Image analysis - scoring polarity establishment under cue/dosage perturbations

To assess polarity establishment and PAR-2 domain formation (Figure 7), zygotes were scored as follows: ‘Domains’, where a PAR-2 domain is visibly formed at NEBD (or early mitosis); ‘Unclear’, where no membrane domain is achieved, but where there appears to be marginal PAR-2 cortical enrichment; and ‘No domains,’ for embryos that have clearly failed to break symmetry. In the case of *spd-5* embryos, domain-containing embryos were subdivided into ‘Monopolar’ and ‘Bipolar.’

### Image analysis - measurement of membrane:cytoplasm signal ratio

To gauge cortical loading of PAR-2 in embryos at different stages of mitosis (see Figure 7H), analysis was done as follows: an intensity profile was obtained by drawing a 10-pixel-wide (2.6 µm) line perpendicular to PAR-2 domain (bisecting the center of domain), from inside to outside of embryo. With this linescan, a membrane signal was calculated by averaging the peak 5 pixels of the profile (to cover the approximate membrane thickness). In parallel, to obtain a cytoplasmic signal, mean pixel intensity was measured within a 900-pixel rectangular ROI drawn in the embryo interior (excluding nuclear/spindle area). These two values were then used to calculate a membrane:cytoplasm ratio. This analysis was done using simple, custom FIJI and Matlab scripts to aid automation.

### Image analysis - measurement of 2-cell size asymmetry

Area of AB and P1 blastomeres was measured on midplane cross-sections of 2-cell embryos, using manually-defined ROIs or through semi-automated segmentation. The size asymmetry was then calculated by dividing the area of AB by the area of the whole embryo. Analysis was performed with custom code in FIJI (see Figures 3A, 3F-H, S2G-H).

### Image analysis - spindle dynamics

Time series of mitotic zygotes were acquired using DIC microscopy, where samples were imaged from NEBD through telophase at 15 s intervals. Centrosome coordinates were tracked manually using semi-automated code in FIJI. Spindle positioning was then determined by measuring the distance of the anterior centrosome or the posterior centrosome to the anterior-most point of the embryo (La and Lp, respectively - see Figure 5A). Final centrosome positioning was defined at telophase as shown in Figure 5B (fLa and fLp for anterior and posterior centrosomes respectively). To monitor transverse oscillations (as seen in Figure 5C-D), centrosome positioning was measured relatively to the nearest point on the A-P axis (namely, Da for anterior and Dp for posterior). Oscillation index Oi equates to the standard deviation of the set of centrosome positions (relative to A-P axis) measured from late prometaphase through telophase.

### Image analysis - cytokinesis asynchrony in AB and P1 blastomeres

The time lag between AB and P1 divisions was measured in DIC timeseries of live embryos. More specifically, asynchrony was defined as the period between the furrow ingression of both blastomeres (t_AB_ and t_P1_ - see illustration in Figure 3B). Furrowing was timed at the point of cortical indentation immediately preceding membrane ‘folding’.

### Image analysis - PAR asymmetry and cytoplasmic ASI

For cortical PAR asymmetry, we define a signal normalized PAR asymmetry according to the following formula, which takes into account combined aPAR and pPAR signal:

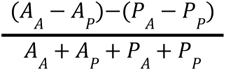

Here, *A*_*A*_ and *A*_*A*_ are the concentrations of aPAR proteins in the anterior and posterior, respectively, and *A*_*P*_ and *A*_*P*_ are the corresponding concentrations of pPAR proteins. Note all concentrations are normalized to the peak concentrations achieved in wild-type embryos. Effectively, this measure defines integrated PAR asymmetry as the sum of the differences in aPAR and pPAR proteins at the two poles, normalized to total signal.

Asymmetry of PAR-1 and MEX-5 patterning was measured in midplane images of zygotes expressing the relevant FP-tagged gene at NEBD. Semi-circle ROIs were drawn on opposite sides of the embryo, and mean pixel intensity for anterior (A) or posterior (P) were retrieved accordingly. ASI was then defined as <u>^*A*^</u> 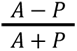 or 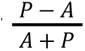 for MEX-5 and PAR-1 respectively, such that ASI is always positive. For quantifying the response of the PAR-1 gradient to PAR-1 depletion, the gradient magnitude (ΔPAR-1) is reported as the absolute concentration difference (P-A) to capture the change in magnitude of the gradient upon PAR-1 dosage reduction. As the relative contributions of membrane and cytoplasmic populations of PAR-1 towards MEX-5 asymmetry are unclear, the semi-circular ROIs were chosen to include both membrane and cytoplasm signals.

### Statistics and formal analysis

Dosage versus phenotypic variance plots (Figure 3) were generated using a moving-window method. For each dataset, a Gaussian weighting function (half-width = 0.1) was moved along the dosage axis, and Gaussian-weighted mean dosage was plotted against Gaussian-weighted phenotypic variance. 95% confidence intervals were determined by bootstrapping.

Unless otherwise specified, statistical analysis was performed in Prism (Graphpad).

## Contributions

Conceptualization: N.T.L.R., N.W.G.; Methodology: N.T.L.R., T.B.; Formal analysis: N.T.L.R., T.B., K.N., N.W.G.; Investigation: N.T.L.R., T.B., K.N., N.W.G.; Resources: N.T.L.R., T.B., N.H.; Writing – original draft preparation: N.T.L.R., N.W.G.; Writing – review and editing: All Authors Supervision: N.W.G.; Project administration: N.W.G.; Funding acquisition: N.W.G.

## Acknowledgements

We thank Buzz Baum, Michalis Barkoulas, and the Goehring Lab for comments on the manuscript. Strains and/or reagents were graciously provided by Dan Dickinson, Ken Kemphues, and Geraldine Seydoux. Additional strains were provided by the Caenorhabditis Genome Center (CGC), which is funded by NIH Office of Research Infrastructure Programs (P40 OD010440), and the Mitani Lab via the National Bio-Resource Project of the MEXT, Japan. This work was supported by the Francis Crick Institute, which receives its core funding from Cancer Research UK (FC001086), the UK Medical Research Council (FC001086), and the Wellcome Trust (FC001086).

For the purpose of Open Access, the author has applied a CC BY public copyright license to any Author Accepted Manuscript version arising from this submission.

---

**Supplemental Figure S1.**
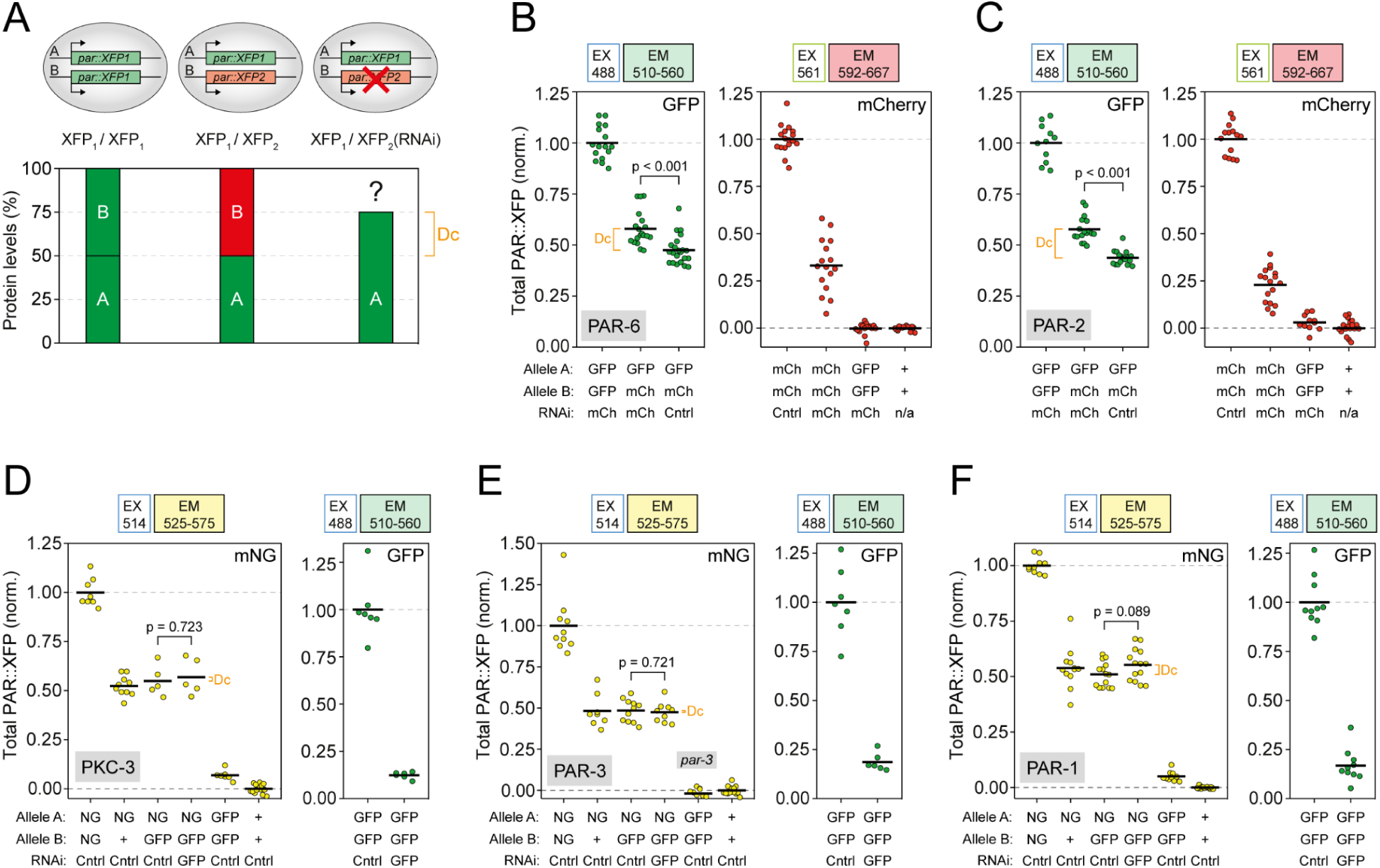
Lack of homeostatic dosage compensation of *par* gene expression in heterozygous animals. **(A)** Schematic for dosage compensation assay using allele-specific RNAi-depletion. **(B-C)** PAR-6::GFP (B) and GFP::PAR-2 (C) levels for *gfp/gfp*, *gfp/mCherry*, and *gfp/mCherry(RNAi)* genotypes (left) together with controls for the depletion of mCherry-tagged alleles by RNAi (right). Note that compensation, if present, should be manifest as a difference in GFP levels in *gfp/mCherry* embryos ± *mCherry(RNAi)*. *gfp/gfp* and *+/+* embryos are shown to control for bleedthrough into the mCherry channel and to confirm zero point, respectively. **(D-F)** mNG::PKC-3 (D), mNG::PAR-3 (E), PAR-1::mNG (F) levels for the indicated genotypes (left) together with controls for depletion of GFP-tagged alleles by RNAi (right). Note that compensation, if present, should be manifest as a difference in mNG levels in *mNG/gfp* embryos ± *gfp(RNAi)*. Relevant samples were compared statistically using an unpaired t test. *gfp/gfp* and +/+ embryos are shown as controls for specificity of mNG excitation and to confirm zero point, respectively.

**Supplemental Figure S2.**
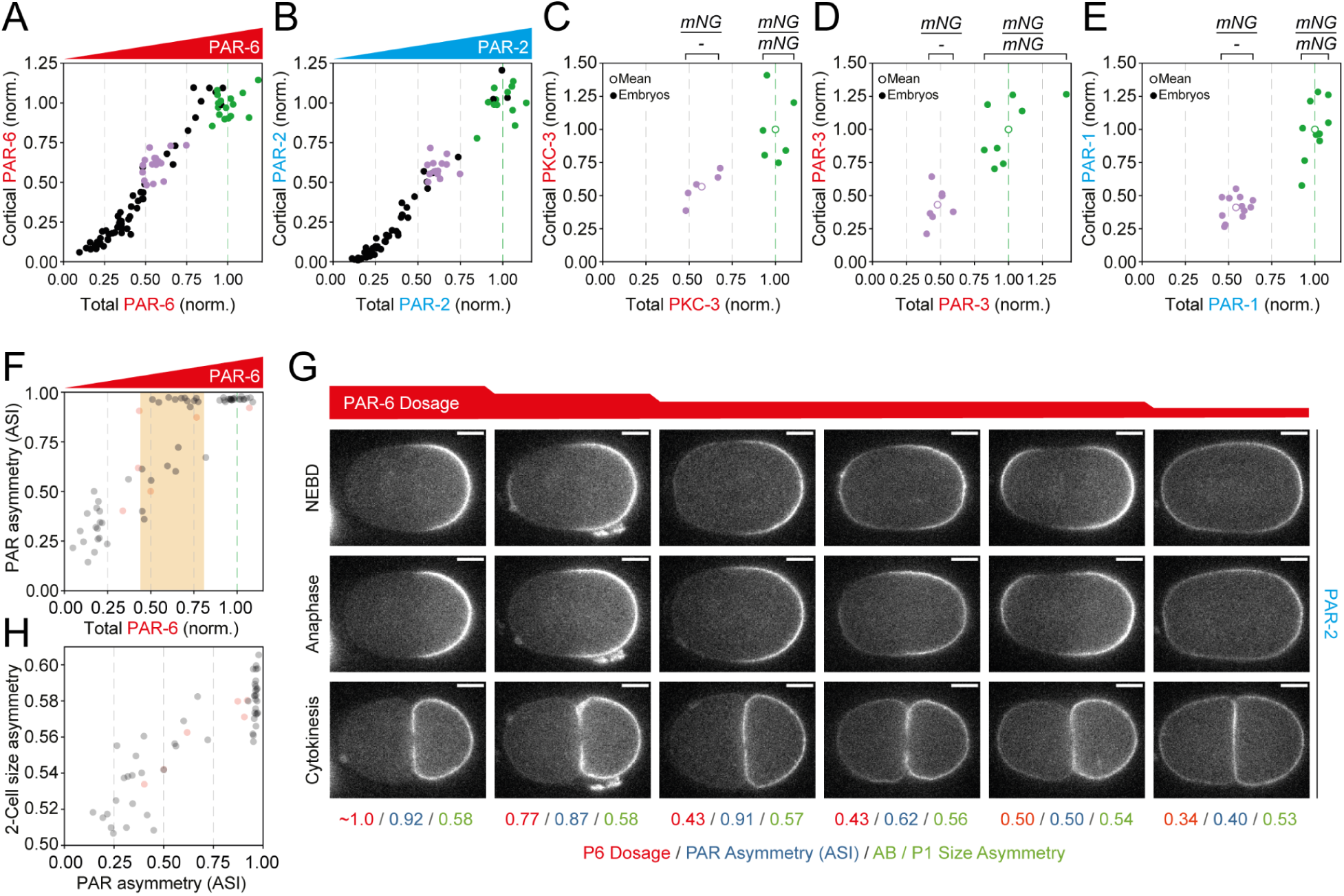
Polarity is robust despite sensitivity of the PAR network to dosage changes. **(A-B)** Cortical concentrations of PAR-6::GFP(A, NWG0119) and GFP::PAR-2(B, NWG0167) decline as a function of total dosage.**(C-E)** Cortical concentrations of PKC-3(C), PAR-3(D), and PAR-1(E) are reduced ∼50% in heterozygous (*mNG/-*) embryos relative to homozygous (*mNG/mNG*) controls. Colored data points in (A-E) indicate homozygous (*xfp/xfp*, green) and heterozygous (*xfp/-*, purple) embryos. Note heterozygous conditions in (A-E) were *par-X(xfp_1_/xfp_2_(RNAi))* as in Figure 2D. **(F)** PAR asymmetry (ASI) as a function of total PAR-6 dosage. Note bimodal behavior with one population in which asymmetry is constant as a function of dosage (PAR-6 > 0.5, PAR asymmetry ∼ 1.0) and a second in which asymmetry declines with dosage (PAR-6 < 0.75, PAR asymmetry < 0.75). **(G)** Example embryos showing PAR-2 localization at NEBD, anaphase and cytokinesis for differing PAR-6 dosage. Note dosage, PAR asymmetry, and size asymmetry are shown below each embryo. Note embryos shown in (G) are depicted as red data points in (F, H). **(H)** Two-cell size asymmetry as a function of PAR asymmetry. Scale bars, 10µm.

**Table S1.**
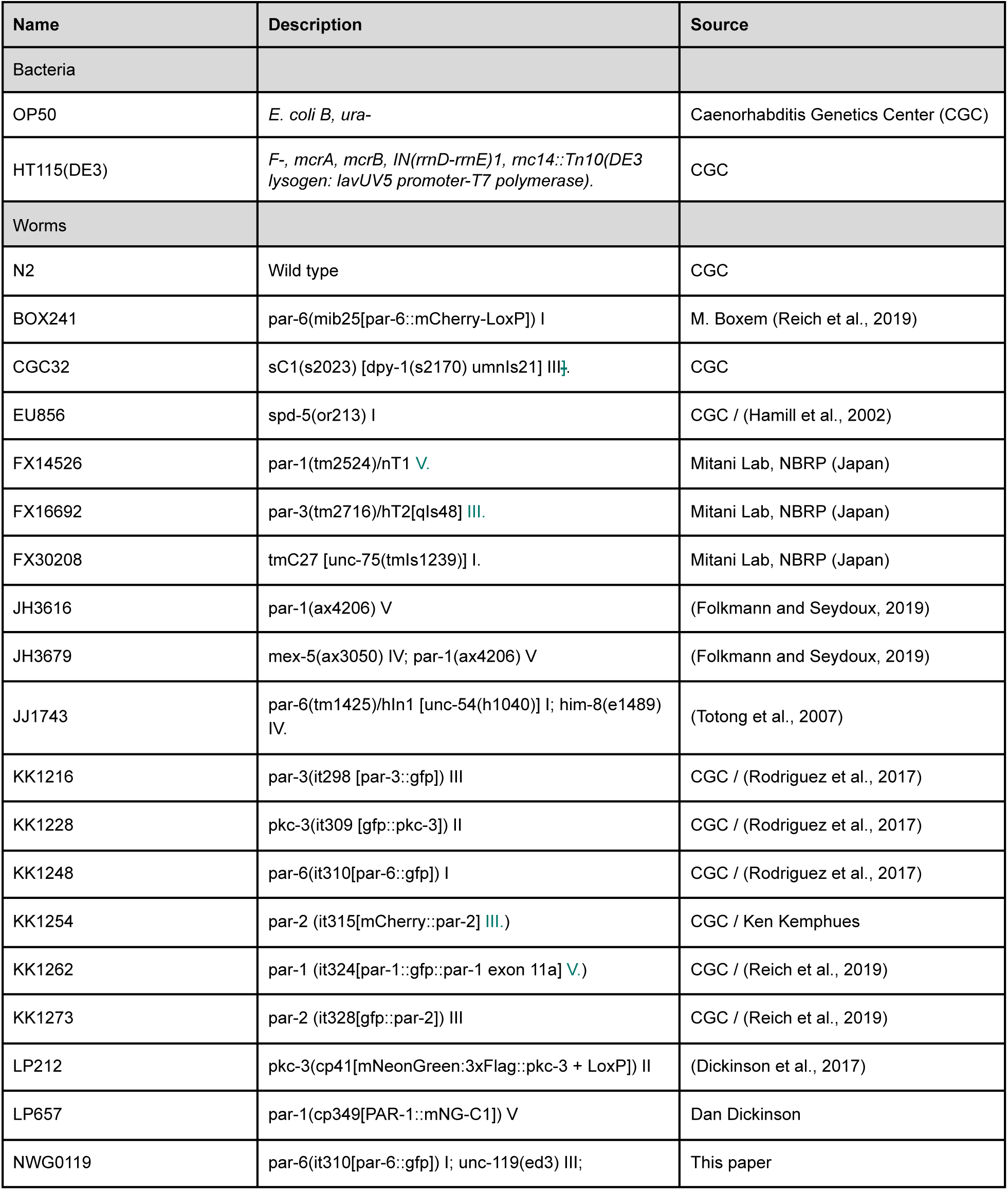
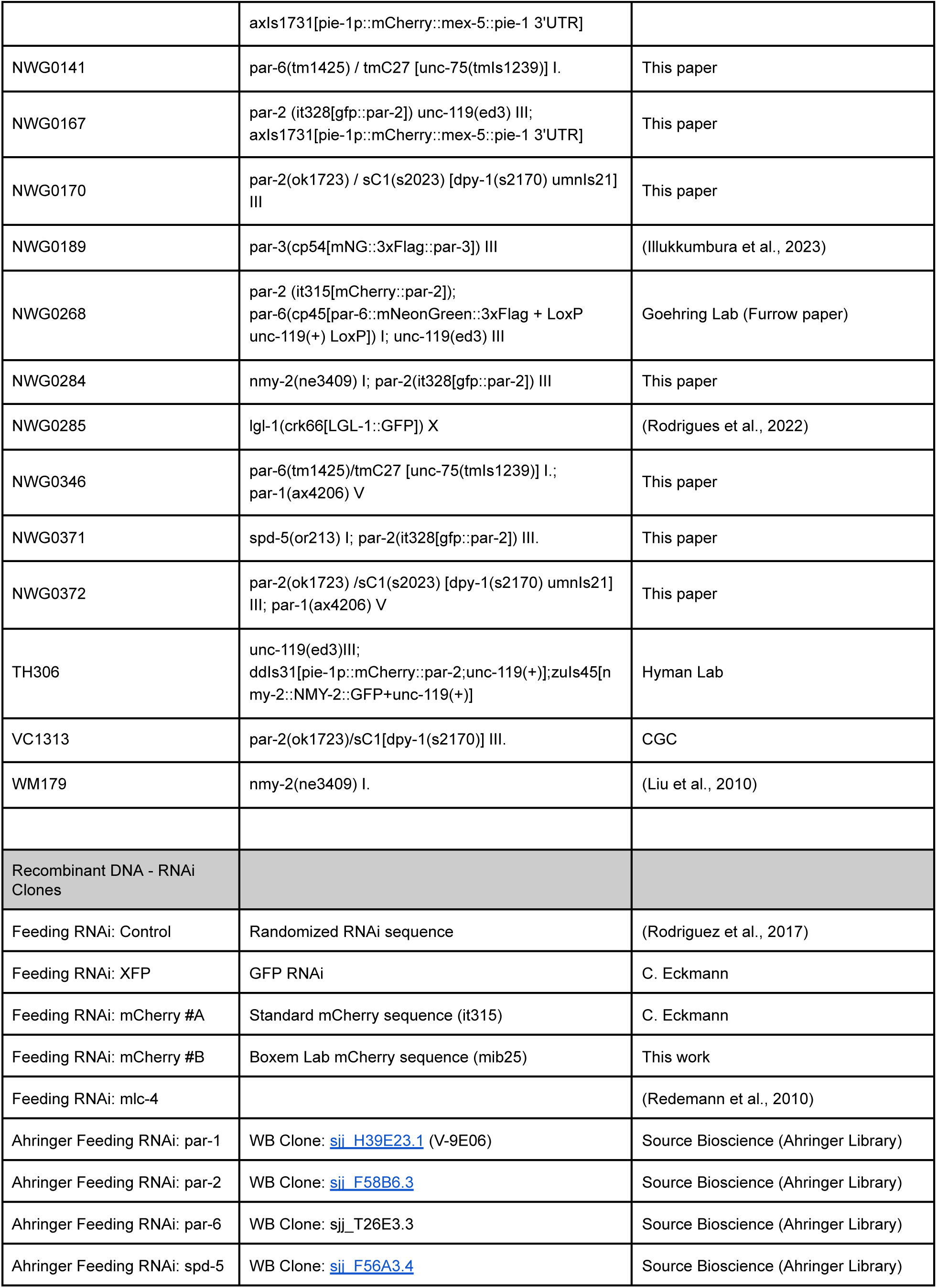
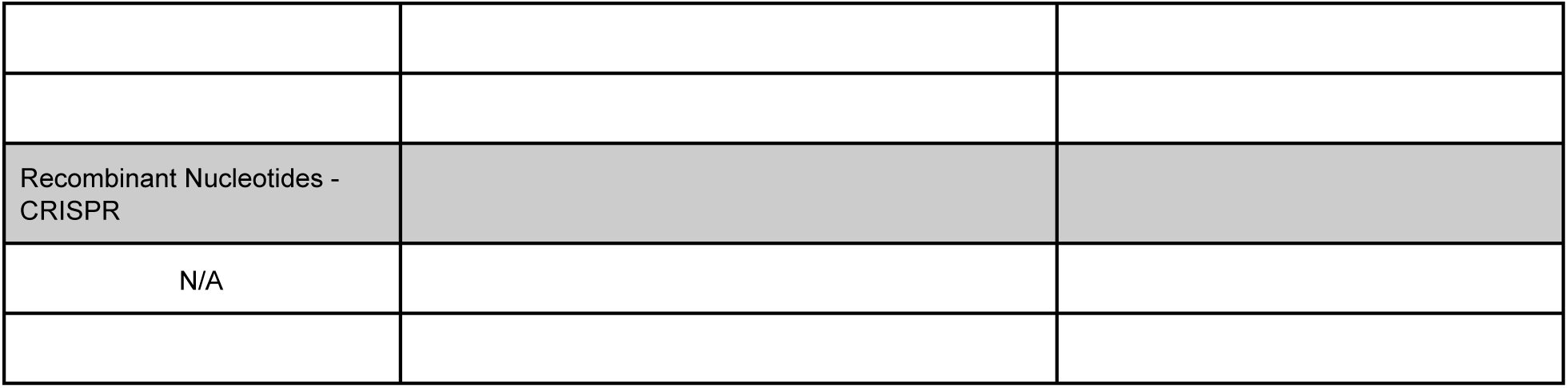
Strains and reagents.

